# Testing for ancient selection using cross-population allele frequency differentiation

**DOI:** 10.1101/017566

**Authors:** Fernando Racimo

## Abstract

A powerful way to detect selection in a population is by modeling local allele frequency changes in a particular region of the genome under scenarios of selection and neutrality, and finding which model is most compatible with the data. Chen et al. [2010] developed a composite likelihood method called XPCLR that uses an outgroup population to detect departures from neutrality which could be compatible with hard or soft sweeps, at linked sites near a beneficial allele. However, this method is most sensitive to recent selection and may miss selective events that happened a long time ago. To overcome this, we developed an extension of XP-CLR that jointly models the behavior of a selected allele in a three-population tree. Our method - called 3P-CLR - outperforms XP-CLR when testing for selection that occurred before two populations split from each other, and can distinguish between those events and events that occurred specifically in each of the populations after the split. We applied our new test to population genomic data from the 1000 Genomes Project, to search for selective sweeps that occurred before the split of Yoruba and Eurasians, but after their split from Neanderthals, and that could have led to the spread of modern-human-specific phenotypes. We also searched for sweep events that occurred in East Asians, Europeans and the ancestors of both populations, after their split from Yoruba. In both cases, we are able to confirm a number of regions identified by previous methods, and find several new candidates for selection in recent and ancient times. For some of these, we also find suggestive functional mutations that may have driven the selective events.

## 2 Introduction

Genetic hitchhiking will distort allele frequency patterns at regions of the genome linked to a beneficial allele that is rising in frequency [Smith and Haigh, 1974]. This is known as a selective sweep. If the sweep is restricted to a particular population and does not affect other closely related populations, one can detect such an event by looking for extreme patterns of localized population differentation, like high values of *F_st_* at a specific locus [Lewontin and Krakauer, 1973]. This and other related statistics have been used to scan the genomes of present-day humans from different populations, so as to detect signals of recent positive selection [Akey et al., 2002, Oleksyk et al., 2008, Weir et al., 2005, Yi et al., 2010].

Once it became possible to sequence entire genomes of archaic humans (like Neanderthals) [Green et al., 2010, Meyer et al., 2012, Prüfer et al., 2014], researchers also began to search for selective sweeps that occurred in the ancestral population of all present-day humans. For example, Green et al. [2010] searched for genomic regions with a depletion of derived alleles in a low-coverage Neanderthal genome, relative to what would be expected given the derived allele frequency in present-day humans. This is a pattern that would be consistent with a sweep in present-day humans. Later on, Prüfer et al. [2014] developed a hidden Markov model (HMM) that could identify regions where Neanderthals fall outside of all present-day human variation (also called "external regions"), and are therefore likely to have been affected by ancient sweeps in early modern humans. They applied their method to a high-coverage Neanderthal genome. Then, they ranked these regions by their genetic length, to find segments that were extremely long, and therefore highly compatible with a selective sweep. Finally, Racimo et al. [2014] used summary statistics calculated in the neighborhood of sites that were ancestral in archaic humans but fixed derived in all or almost all present-day humans, to test if any of these sites could be compatible with a selective sweep model. While these methods harnessed different summaries of the patterns of differentiation left by sweeps, they did not attempt to explicitly model the process by which these patterns are generated over time.

Chen et al. [2010] developed a method called XP-CLR, which is designed to test for selection in one population after its split from a second, outgroup, population *t_AB_* generations ago. It does so by modeling the evolutionary trajectory of an allele under linked selection and under neutrality, and then comparing the likelihood of the data for each of the two models. The method detects local allele frequency differences that are compatible with the linked selection model [Smith and Haigh, 1974], along windows of the genome.

XP-CLR is a powerful test for detecting selective events restricted to one population. However, it provides little information about when these events happened, as it models all sweeps as if they had immediately occurred in the present generation. Additionally, if one is interested in selective sweeps that took place before two populations *a* and *b* split from each other, one would have to run XP-CLR separately on each population, with a third outgroup population c that split from the ancestor of *a* and *b t_ABC_* generations ago (with *t_ABC_* > *t_AB_*). Then, one would need to check that the signal of selection appears in both tests. This may miss important information about correlated allele frequency changes shared by *a* and *b*, but not by *c*, limiting the power to detect ancient events.

To overcome this, we developed an extension of XP-CLR that jointly models the behavior of an allele in all 3 populations, to detect selective events that occurred before or after the closest two populations split from each other. Below we briefly review the modeling framework of XP-CLR and describe our new test, which we call 3P-CLR. In the Results, we show this method outperforms XP-CLR when testing for selection that occurred before the split of two populations, and can distinguish between those events and events that occurred after the split, unlike XP-CLR. We then apply the method to population genomic data from the 1000 Genomes Project [Abecasis et al., 2012], to search for selective sweep events that occurred before the split of Yoruba and Eurasians, but after their split from Neanderthals. We also use it to search for selective sweeps that occurred in the Eurasian ancestral population, and to distinguish those from events that occurred specifically in East Asians or specifically in Europeans.

## 3 Materials and Methods

### 3.1 XP-CLR

First, we review the procedure used by XP-CLR to model the evolution of allele frequency changes of two populations *a* and *b* that split from each other *t_AB_* generations ago (Figure 1.A). For neutral SNPs, Chen et al. [2010] use an approximation to the Wright-Fisher diffusion dynamics [Nicholson et al., 2002]. Namely, the frequency of a SNP in a population *a* (*p_A_*) in the present is treated as a random variable governed by a normal distribution with mean equal to the frequency in the ancestral population (*β*) and variance proportional to the drift time *ω* from the ancestral to the present population:

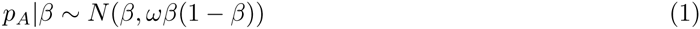

**Figure 1.**
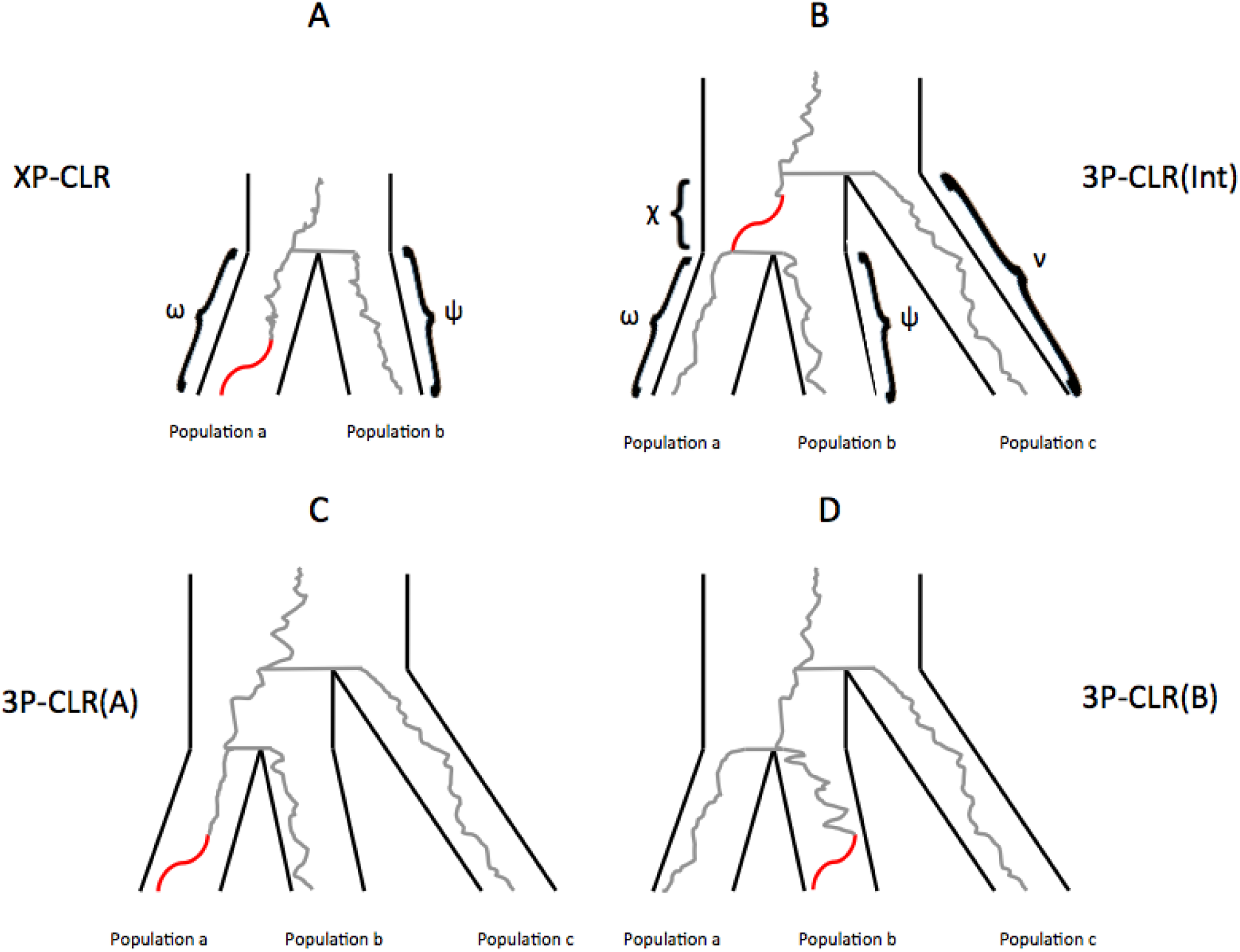
Schematic tree of selective sweeps detected by XP-CLR and 3P-CLR. While XP-CLR can only use two populations (an outgroup and a test) to detect selection (panel A), 3P-CLR can detect selection in the ancestral branch of two populations (3P-CLR(Int), panel B) or on the branches specific to each population (3P-CLR(A) and 3P-CLR(B), panels C and D, respectively). The greek letters denote the known drift times for each branch of the population tree.

where *ω* = *t_AB_/*(*2N_e_)* and *N_e_* is the effective size of population A.

This is a Brownian motion approximation to the Wright-Fisher model, as the drift increment to variance is constant across generations. If a SNP is segregating in both populations - i.e. has not hit the boundaries of fixation or extinction - this process is time-reversible. Thus, one can model the frequency of the SNP in population *a* with a normal distribution having mean equal to the frequency in population *b* and variance proportional to the sum of the drift time (*ω*) between *a* and the ancestral population, and the drift time between b and the ancestral population (*ψ*):

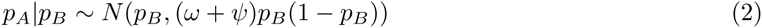

For SNPs that are linked to a beneficial allele that has produced a sweep in population a only, Chen et al. [2010] model the allele as evolving neutrally until the present and then apply a transformation to the normal distribution that depends on the distance to the selected allele r and the strength of selections [Durrett and Schweinsberg, 2004, Fay and Wu, 2000]. Let 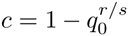 where *q*_0_ is the frequency of the beneficial allele in population *a* before the sweep begins. The frequency of a neutral allele is expected to increase from p to l − *c* + *cp* if the allele is linked to the beneficial allele, and this occurs with probability equal to the frequency of the neutral allele (*p*) before the sweep begins. Otherwise, the frequency of the neutral allele is expected to decrease from *p* to *cp.* This leads to the following transformation of the normal distribution:

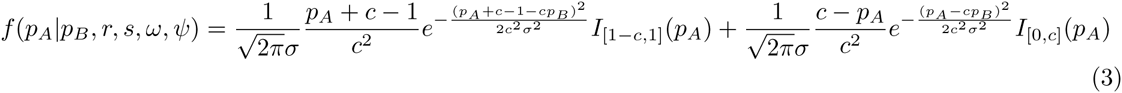

where *ó*^2^ = (*ù* + *ø*)*p_B_*(1 − *p_B_*) and *I*_[_*_x;y_*_]_(*z*) is 1 on the interval [*x*; *y*] and 0 otherwise.

For *s* → 0 or *r* >> *s*, this distribution converges to the neutral case. Let **v** be the vector of all drift times that are relevant to the scenario we are studying. In this case, it will be equal to (*ω*; *ψ*) but in more complex cases below, it may include additional drift times. Let r be the vector of recombination fractions between the beneficial alleles and each of the SNPs within a window of arbitrary size. We can then calculate the product of likelihoods over all k SNPs in that window for either the neutral or the linked selection model, after binomial sampling of alleles from the population frequency, and conditioning on the event that the allele is segregating in the population:

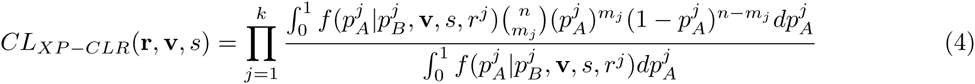

This is a composite likelihood [Lindsay, 1988, Varin et al., 2011], because we are ignoring the correlation in frequencies produced by linkage among SNPs that is not strictly due to proximity to the beneficial SNP. We note that the denominator in the above equation is not explicitly stated in Chen et al. [2010] for ease of notation, but appears in the published online implementation of the method.

Finally, we obtain a composite likelihood ratio statistic *S_XP–CLR_* of the hypothesis of linked selection over the hypothesis of neutrality:

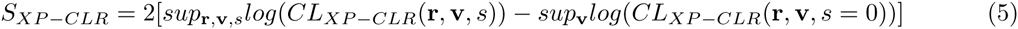

For ease of computation, Chen et al. [2010] assume that r is given (via a recombination map) instead of maximizing the likelihood with respect to it, and we will do so too. Furthermore, they empirically estimate v using *F*_2_ statistics [Patterson et al., 2012] calculated over the whole genome, and assume selection is not strong or frequent enough to affect their genome-wide values. Therefore, the likelihoods in the above equation are only maximized with respect to the selection coefficient, using a grid of coefficients on a logarithmic scale.

### 3.2 3P-CLR

We are interested in the case where a selective event occurred more anciently than the split of two populations (*a* and *b*) from each other, but more recently than their split from a third population *c* (Figure 1.B). We begin by modeling *p_A_* and *p_B_* as evolving from an unknown common ancestral frequency *β*:

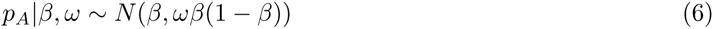

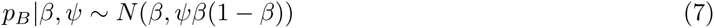

Let *χ* be the drift time separating the most recent common ancestor of *a* and *b* from the most recent common ancestor of *a*, *b* and *c*. Additionally, let *v* be the drift time separating population *c* in the present from the most recent common ancestor of *a*, *b* and *c*. Given these parameters, we can treat *β* as an additional random variable that either evolves neutrally or is linked to a selected allele that swept immediately more anciently than the split of *a* and *b*. In both cases, the distribution of *β* will depend on the frequency of the allele in population *c* (*p_C_*) in the present. In the neutral case:

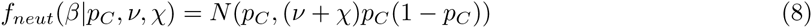

In the linked selection case:

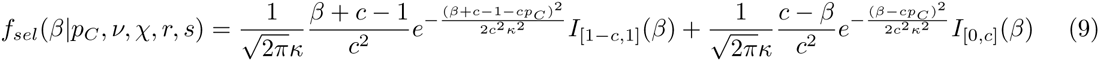

where *κ*^2^ = (*ν* + *χ*)*p_C_*(1 –*p_C_*)

The frequencies in *a* and *b* given the frequency in *c* can be obtained by integrating *β* out. This leads to a density function that models selection in the ancestral population of *a* and *b*.

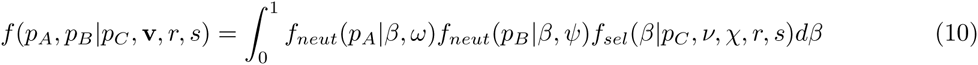

Additionally, formula 10 can be modified to test for selection that occurred specifically in one of the terminal branches that lead to *a* or *b* (Figures 1.C and 1.D), rather than in the ancestral population of *a* and *b*. For example, the density of frequencies for a scenario of selection in the branch leading to *a* can be written as:

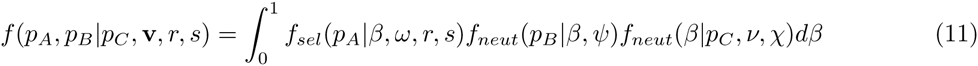

We will henceforth refer to the version of 3P-CLR that is tailored to detect selection in the internal branch that is ancestral to *a* and *b* as 3P-CLR(Int). In turn, the versions of 3P-CLR that are designed to detect selection in each of the daughter populations *a* and *b* will be designated as 3P-CLR(A) and 3P-CLR(B), respectively.

We can now calculate the probability density of specific allele frequencies in populations *a* and *b*, given that we observe *m_C_* derived alleles in a sample of size *n_C_* from population *c*:

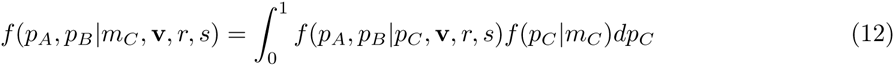

and

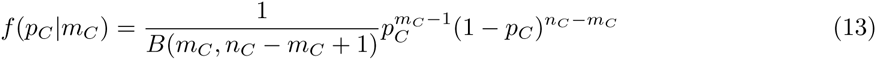

where B(x,y) is the Beta function. We note that formula 13 assumes that the unconditioned density function for the population derived allele frequency *f*(*p_C_*) comes from the neutral infinite-sites model at equilibrium and is therefore equal to the product of a constant and 1/*p_C_* [Ewens, 2012].

Conditioning on the event that the site is segregating in the population, we can then calculate the probability of observing *m_A_* and *m_B_* derived alleles in a sample of size *n_A_* from population *a* and a sample of size *n_B_* from population *b*, respectively, given that we observe *m_C_* derived alleles in a sample of size *n_C_* from population *c*, using binomial sampling:

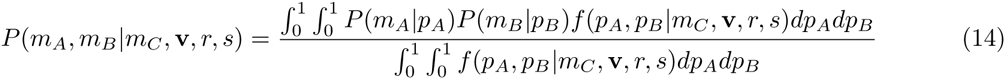

where

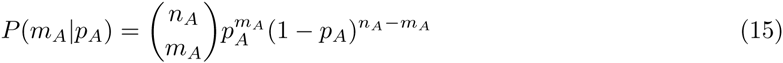

and

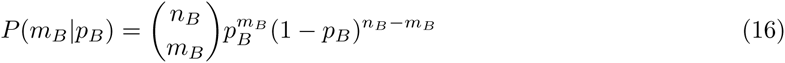

This allows us to calculate a composite likelihood of the derived allele counts in *a* and *b* given the derived allele counts in *c*:

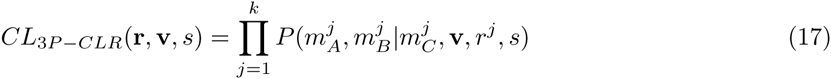

As before, we can use this composite likelihood to produce a composite likelihood ratio statistic that can be calculated over regions of the genome to test the hypothesis of linked selection centered on a particular locus against the hypothesis of neutrality. Due to computational costs in numerical integration, we skip the sampling step for population *c* (formula 13) in our implementation of 3P-CLR. In other words, we assume *p_C_* = *m_C_/n_C_*, but this is also assumed in XP-CLR when computing its corresponding outgroup frequency. To perform the numerical integrations, we used the package Cubature (v.1.0.2). We implemented our method in a freely available C++ program that can be downloaded from here:

https://github.com/ferracimo

The program requires the neutral drift parameters *α*, *β* and (*ν* + *χ*) to be specified as input. These can be obtained using *F*_3_ statistics [Felsenstein, 1981, Patterson et al., 2012], which have previously been implemented in programs like MixMapper [Lipson et al., 2013]. For example, *α* can be obtained via *F*_3_(*A*; *B*; *C*), while (*ν* + *χ*) can be obtained via *F*_3_(*C*; *A*; *B*). When computing *F*_3_ statistics, we use only sites where population C is polymorphic, and so we correct for this ascertainment in the calculation. Another way of calculating these drift times is via ∂*a*∂*i* [Gutenkunst et al., 2009]. Focusing on two populations at a time, we can fix one population’s size and allow the split time and the other population’s size to be estimated by the program, in this case using all polymorphic sites, regardless of which population they are segregating in. We then obtain the two drift times by scaling the inferred split time by the two different population sizes. We provide scripts in our github page for the user to obtain these drift parameters using both of the above ways.

## 4 Results

### 4.1 Simulations

We generated simulations in SLiM [Messer, 2013] to test the performance of XP-CLR and 3P-CLR in a three-population scenario. We first focused on the performance of 3P-CLR(Int) in detecting ancient selective events that occurred in the ancestral branch of two sister populations. We assumed that the population history had been correctly estimated (i.e. the drift parameters and population topology were known). First, we simulated scenarios in which a beneficial mutation arose in the ancestor of populations *a* and *b*, before their split from each other but after their split from *c* (Table 1). Although both XP-CLR and 3P-CLR are sensitive to partial or soft sweeps (as they do not rely on extended patterns of haplotype homozygosity [Chen et al., 2010]), we required the beneficial allele to have fixed before the split (at time *t_ab_*) to ensure that the allele had not been lost by then, and also to ensure that the sweep was restricted to the internal branch of the tree. We fixed the effective size of all three populations at *N_e_* = 10,000. Each simulation consisted in a 5 cM region and the beneficial mutation occurred in the center of this region. The mutation rate was set at 2.5 * 10^−8^ per generation and the recombination rate between adjacent nucleotides was set at 10^−8^ per generation.

**Table 1.** Description of models tested. All times are in generations. Selection in the "ancestral population" refers to a selective sweep where the beneficial mutation and fixation occurred before the split time of the two most closely related populations. Selection in "daughter population *a*" refers to a selective sweep that occurred in one of the two most closely related populations (a), after their split from each other. *t_AB_*: split time (in generations ago) of populations *a* and *b*. *t_ABC_*: split time of population *c* and the ancestral population of *a* and *b*. *t_M_*: time at which the selected mutation is introduced. *s*: selection coefficient. *N_e_*: effective population size.

| Model | Population where selection occurred | $t_{AB}$ | $t_{ABC}$ | $t_M$ | *s* | $N_e$ |
| --- | --- | --- | --- | --- | --- | --- |
| A | Ancestral population | 500 | 2,000 | 1,800 | 0.1 | 10,000 |
| B | Ancestral population | 1,000 | 4,000 | 2,500 | 0.1 | 10,000 |
| C | Ancestral population | 2,000 | 4,000 | 3,500 | 0.1 | 10,000 |
| D | Ancestral population | 3,000 | 8,000 | 5,000 | 0.1 | 10,000 |
| E | Ancestral population | 2,000 | 16,000 | 8,000 | 0.1 | 10,000 |
| F | Ancestral population | 4,000 | 16,000 | 8,000 | 0.1 | 10,000 |
| I | Daughter population *a* | 2,000 | 4,000 | 1,000 | 0.1 | 10,000 |
| J | Daughter population *a* | 3,000 | 8,000 | 2,000 | 0.1 | 10,000 |

**Table 2.**
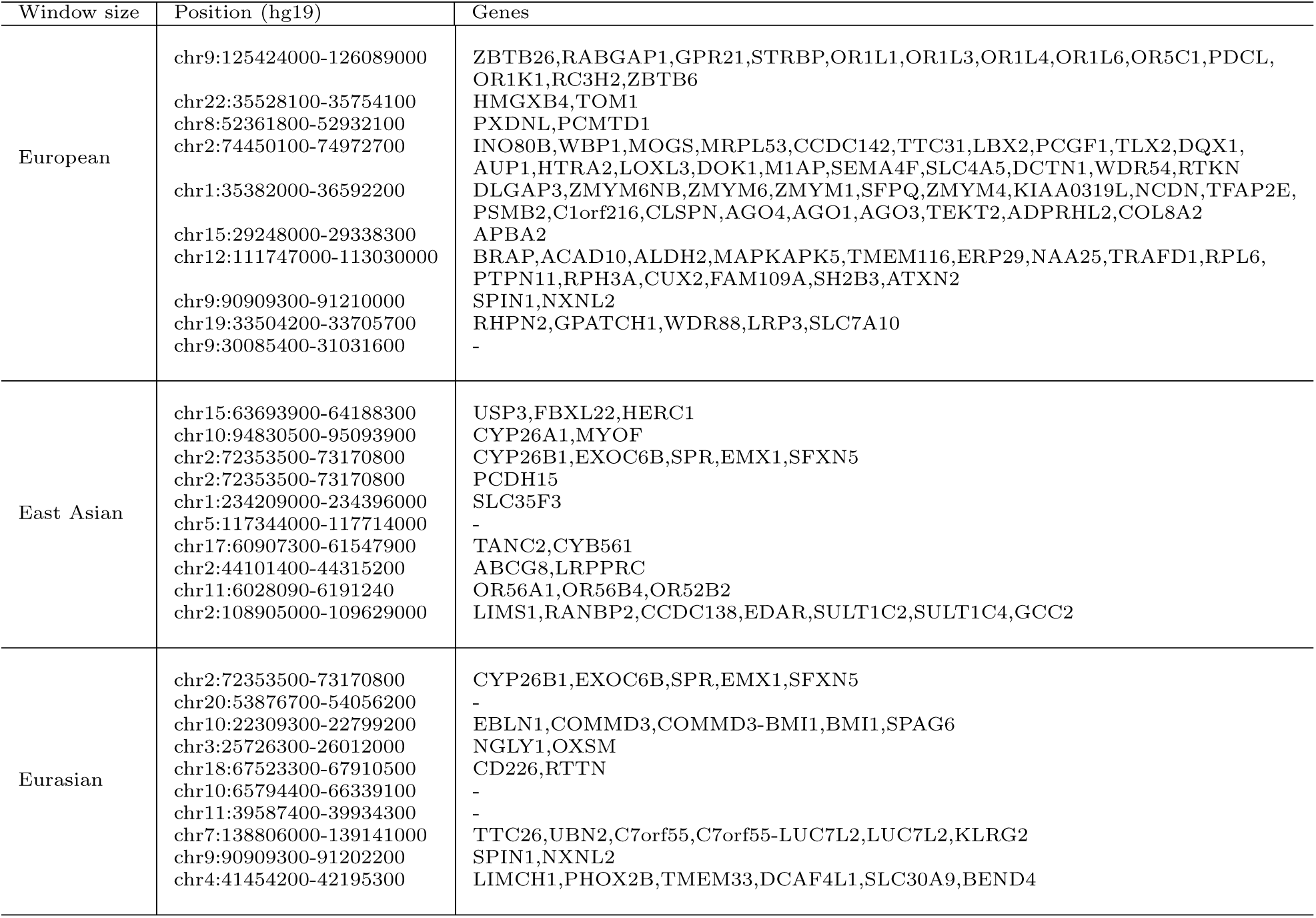
Genes from top 10 candidate regions for each of the branches on which 3P-CLR was run for the Eurasian population tree. All positions were rounded to the nearest 100 bp. Windows were merged together if the central SNPs that define them were contiguous.

To make a fair comparison to 3P-CLR(Int), and given that XP-CLR is a two-population test, we applied XP-CLR in two ways. First, we pretended population *b* was not sampled, and so the "test" panel consisted of individuals from *a* only, while the "outgroup" consisted of individuals from *c*. In the second implementation (which we call "XP-CLR-avg"), we used the same outgroup panel, but pooled the individulas from *a* and *b* into a single panel, and this pooled panel was the "test". The window size was set at 0.5 cM and the number of SNPs sampled between each window’s central SNP was set at 600 (this number is large because it includes SNPs that are not segregating in the outgroup, which are later discarded). To speed up computation, and because we are largely interested in comparing the relative performance of the three tests under different scenarios, we used only 20 randomly chosen SNPs per window in all tests. We note, however, that the performance of all of these tests can be improved by using more SNPs per window.

Figure 2 shows receiver operating characteristic (ROC) curves comparing the sensitivity and specificity of 3P-CLR(Int), 3P-CLR(A), XP-CLR and XP-CLR-avg in the first six demographic scenarios described in Table 1. Each ROC curve was made from 100 simulations under selection (with *s* = 0.1 for the central mutation) and 100 simulations under neutrality (with *s* = 0 and no fixation required). In each simulation, 100 haploid individuals (or 50 diploids) were sampled from population *a*, 100 individuals from population *b* and 100 individuals from the outgroup population *c*. For each simulation, we took the maximum value at a region in the neighborhood of the central mutation (+/− 0.5 cM) and used those values to compute ROC curves under the two models.

**Figure 2.**
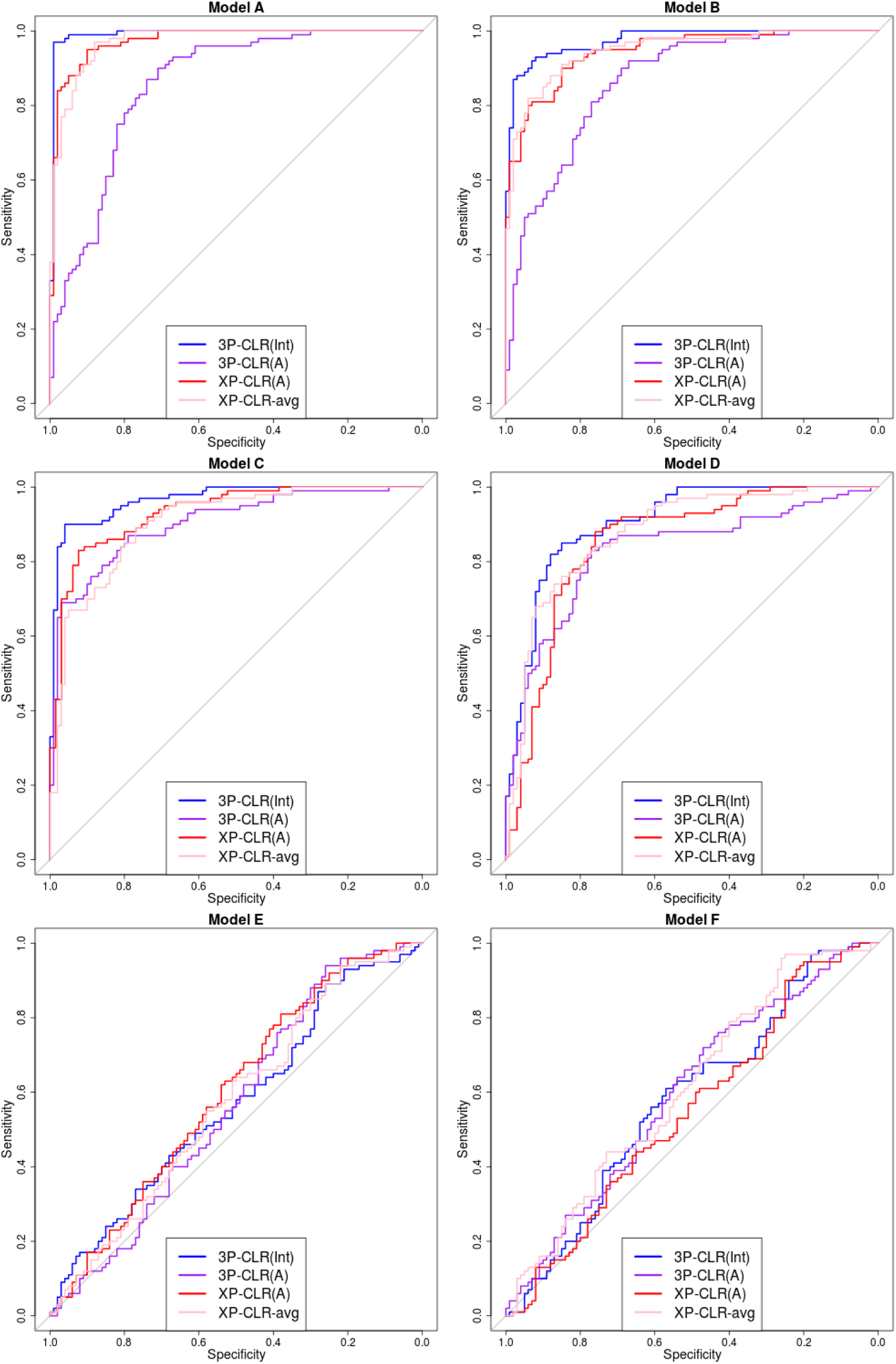
ROC curves for performance of 3P-CLR(Int), 3P-CLR(A) and two variants of XP-CLR in detecting selective sweeps that occurred before the split of two populations *a* and *b*, under different demographic models. In this case, the outgroup panel from population *c* contained 100 haploid genomes. The two sister population panels (from *a* and *b*) also have 100 haploid genomes each.

When the split times are recent or moderately ancient (models A to D), 3P-CLR(Int) outperforms the two versions of XP-CLR. Furthermore, 3P-CLR(A) is the test that is least sensitive to selection in the internal branch as it is only meant to detect selection in the terminal branch leading to population *a*. When the split times are very ancient (models E and F), none of the tests perform well. The root mean squared error (RMSE) of the genetic distance between the true selected site and the highest scored window is comparable across tests in all six scenarios (Figure S5). 3P-CLR(Int) is the best test at finding the true location of the selected site in almost all demographic scenarios. We observe that we lose almost all power if we simulate demographic scenarios where the population size is 10 times smaller (*N_e_* = 1,000) (Figure S1). Additionally, we observe that the power and specificity of 3P-CLR decrease as the selection coefficient decreases (Figure S2).

We also simulated a situation in which only a few individuals are sequenced from the outgroup, while large numbers of sequences are available from the tests. Figures S3 and S6 show the ROC curves and RMSE plots, respectively, for a scenario in which 100 individuals were sampled from the test populations but only 10 individuals (5 diploids) were sampled from the outgroup. Unsurprisingly, all tests have less power to detect selection when the split times and the selection events are recent to moderately ancient (models A-D). Interestingly though, when the split times and the selective events are very ancient (models E-F), both 3P-CLR and XP-CLR perform better when using a small ougroup panel (Figure S3) than when using a large outgroup panel (Figure 2). This is due to the Brownian motion approximation that these methods utilize. Under the Wright-Fisher model, the drift increment at generation t is proportional to p(t)*(1-p(t)), where p(t) is the derived allele frequency. The derivative of this function gets smaller the closer p(t) is to 0.5 (and is exactly 0 at that point). Small outgroup panels serve to filter out loci with allele frequencies far from 0.5, and so small changes in allele frequency will not affect the drift increment much, making Brownian motion a good approximation to the Wright-Fisher model. Indeed, when running 3P-ClR(Int) in a demographic scenario with very ancient split times (Model E) and a large outgroup panel (100 sequences) but only restricting to sites that are at intermediate frequencies in the outgroup (25% ≤ *m_C_*/*n_C_* ≤ 75%), we find that performance is much improved relative to the case when we use all sites that are segregating in the outgroup (Figure S4).

Importantly, the usefulness of 3P-CLR(Int) resides not just in its performance at detecting selective sweeps in the ancestral population, but in its specific sensitivity to that particular type of events. Because the test relies on correlated allele frequency differences in both population *a* and population *b* (relative to the outgroup), selective sweeps that are specific to only one of the populations will not lead to high 3P-CLR(Int) scores, but will instead lead to high 3P-CLR(A) scores or 3P-CLR(B) scores, depending on where selection took place. Figure 3 shows ROC curves in two scenarios in which a selective sweep occurred only in population *a* (models I and J in Table 1), using 100 sampled individuals from each of the 3 populations. Here, XP-CLR performs well, but is outperformed by 3P-CLR(A). Furthermore, 3P-CLR(Int) shows almost no sensitivity to the recent sweep. For example, in Model I, at a specificity of 90%, 3P-CLR(A) and XP-CLR(A) have 86% and 80% sensitivity, respectively, while at the same specificity, 3P-CLR(Int) only has 18% sensitivity. One can compare this to the same demographic scenario but with selection occurring in the ancestral population of *a* and *b* (model C, Figure 2), where at 90% specificity, 3P-CLR(A) and XP-CLR(A) have 72% and 84% sensitivity, respectively, while 3P-CLR(Int) has 90% sensitivity. We also observe that 3P-CLR(A) is the best test at finding the true location of the selected site when selection occurs in the terminal branch leading to population *a* (Figure S7).

**Figure 3.**
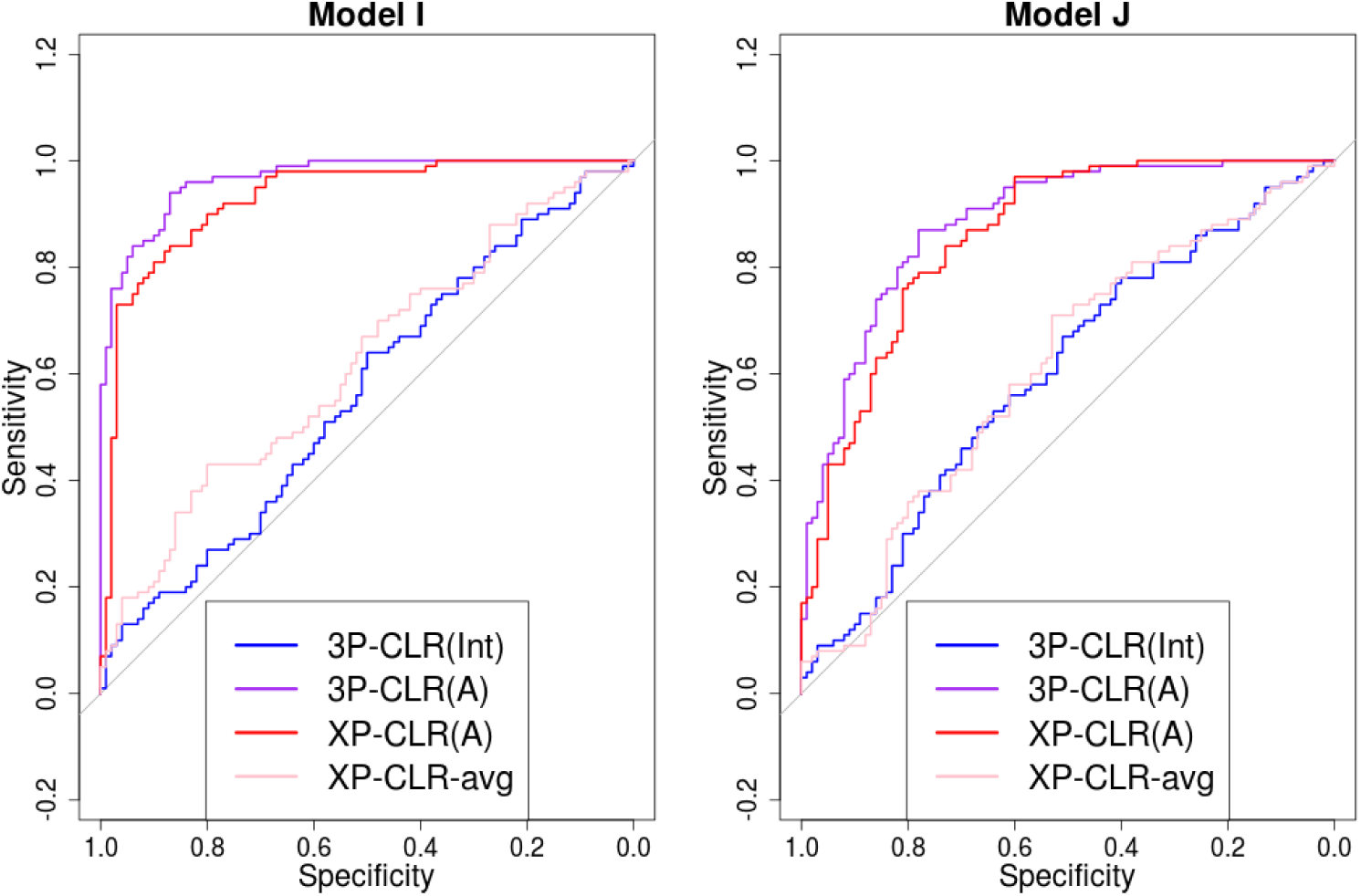
3P-CLR(Int) is tailored to detect selective events that happened before the split *t_ab_*, so it is largely insensitive to sweeps that occurred after the split. ROC curves show performance of 3P-CLR(Int) and two variants of XP-CLR for models where selection occurred in population *a* after its split from *b*.

Finally, we tested the behavior of 3P-CLR under selective scenarios that we did not explicitly model. First, we simulated a selective sweep in the outgroup population. We find that all three types of 3P-CLR statistics (3P-CLR(Int), 3P-CLR(A) and 3P-CLR(B)) are largely insensitive to this type of event, though 3P-CLR(Int) is relatively more sensitive than the other two. Second, we simulated two independent selective sweeps in populations *a* and *b* (convergent evolution). This results in elevated 3P-CLR(A) and 3P-CLR(B) statistics, but 3P-CLR(Int) remains largely insensitive (Figure S8). We note that 3P-CLR should not be used to detect selective events that occurred before the split of all three populations (i.e. before the split of *c* and the ancestor of *a* and *b*), as it relies on allele frequency differences between the populations.

### 4.2 Selection in Eurasians

We first applied 3P-CLR to modern human data from phase 1 of the 1000 Genomes Project [Abecasis et al., 2012]. We used the African-American recombination map [Hinch et al., 2011] to convert physical distances into genetic distances. We focused on Europeans (CEU, FIN, GBR, IBS, TSI) and East Asians (CHB, CHS, JPT) as the two sister populations, using Yoruba (YRI) as the outgroup population (Figure S9.A). We randomly sampled 100 individuals from each population and obtained sample derived allele frequencies every 10 SNPs in the genome. We then calculated likelihood ratio statistics by a sliding window approach, where we sampled a "central SNP" once every 10 SNPs. The central SNP in each window was the candidate beneficial SNP for that window. We set the window size to 0.25 cM, and randomly sampled 100 SNPs from each window, centered around the candidate beneficial SNP. In each window, we calculated 3P-CLR to test for selection at three different branches of the population tree: the terminal branch leading to Europeans (3P-CLR Europe), the terminal branch leading to East Asians (3P-CLR East Asia) and the ancestral branch of Europeans and East Asians (3P-CLR Eurasia). Results are shown in Figure S10. For each scan, we selected the windows in the top 99.9% quantile of scores and merged them together if their corresponding central SNPs were contiguous, effectively resulting in overlapping windows being merged. Tables S1, S2 and S3 show the top hits for Europeans, East Asians and the ancestral Eurasian branch, respectively

We observe several genes that were identified in previous selection scans. In the East Asian branch, one of the top hits is *EDAR.* Figure 4. A shows that this gene appears to be under selection exclusively in this population branch. It codes for a protein involved in hair thickness and incisor tooth morphology [Fujimoto et al., 2008, Kimura et al., 2009], and has been repeatedly identified as a candidate for a sweep in East Asians [Grossman et al., 2010, Sabeti et al., 2007].

**Figure 4.**
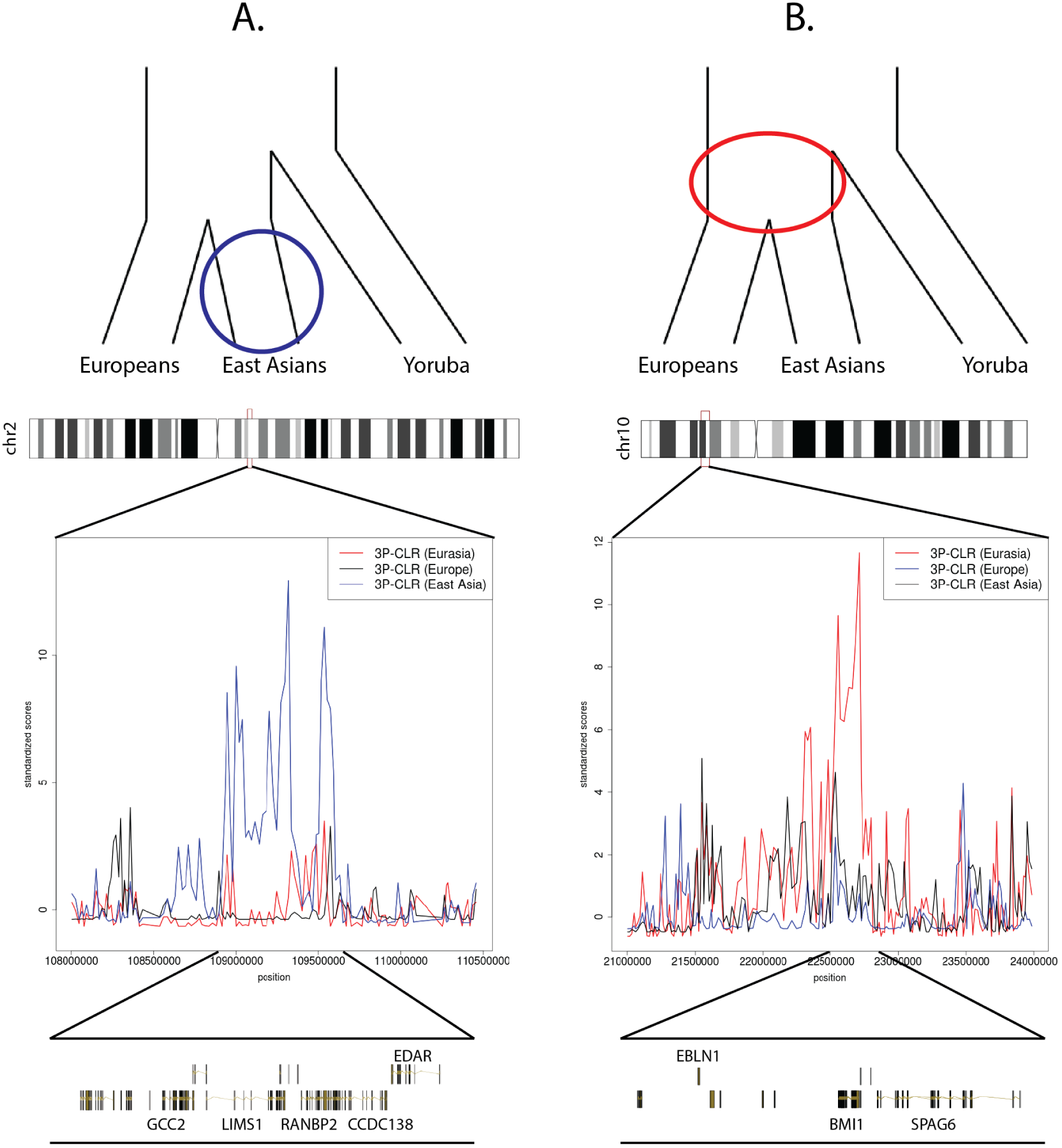
3P-CLR scan of Europeans (black), East Asians (blue) and the ancestral Eurasian population (red) reveals regions under selection in different branches of the population tree. To make a fair comparison, all 3P-CLR scores were standardized by substracting the chromosome-wide mean from each window and dividing the resulting score by the chromosome-wide standard deviation. A) The region containing *EDAR* is a candidate for selection in the East Asian population. B) The region containing genes *SPAG6* and *BMI1* is a candidate for selection in the ancestral population of Europeans and East Asians. The image was built using the GenomeGraphs package in Bioconductor.

Furthermore, 3P-CLR allows us to narrow down on the specific time at which selection for previously found candidates occurred in the history of particular populations. For example, Chen et al. [2010] performed a scan of the genomes of East Asians using XP-CLR with Yoruba as the outgroup, and identified a number of candidate genes [Chen et al., 2010]. 3P-CLR confirms recovers several of their loci when looking specifically at the East Asian branch: *OR56A1, OR56B4, OR52B2, SLC30A9, BBX, EPHB1, ACTN1* and *XKR6.* However, when applied to the ancestral Eurasian branch, 3P-CLR finds some genes that were previously found in the XP-CLR analysis of East Asians, but that are not among the top hits in 3P-CLR applied to the East Asian branch: *COMMD3, BMI1, SPAG6, NGLY1, OXSM, CD226, ABCC12, ABCC11, LONP2, SIAH1, PPARA, PKDREJ, GTSE1, TRMU* and *CELSR1.* This suggests selection in these regions occurred earlier, i.e. before the European-East Asian split. Figure 4.B shows a comparison between the 3P-CLR scores for the three branches in the region containing genes *BMI1* (a proto-oncogene [Siddique and Saleem, 2012]) and *SPAG6* (involved in sperm motility [Sapiro et al., 2002]). Here, the signal of Eurasia-specific selection is evidently stronger than the other two signals. Finally, we also find some candidates from Chen et al. [2010] that appear to be under selection in both the ancestral Eurasian branch and the East Asian daughter branch: *SFXN5, EMX1, SPR* and *CYP26B1.* Interestingly, both CYP26B1 and CYP26A1 are very strong candidates for selection in the East Asian branch. These two genes lie in two different chromosomes, so they are not part of a gene cluster, but they both code for proteins that hydrolize retinoic acid, an important signaling molecule [Topletz et al., 2012, White et al., 2000].

Other selective events that 3P-CLR infers to have occurred in Eurasians include the region containing *HERC2* and *OCA2,* which are major determinants of eye color [Branicki et al., 2009, Eiberg et al., 2008, Han et al., 2008]. There is also evidence that these genes underwent selection more recently in the history of Europeans [Mathieson et al., 2015], which could suggest an extended period of selection - perhaps influenced by migrations between Asia and Europe - or repeated selective events at the same locus.

When running 3P-CLR to look for selection specific to Europe, we find that *TYRP1,* which plays a role in human skin pigmentation [Halaban and Moellmann, 1990], is among the top hits. This gene has been previously found to be under strong selection in Europe [Voight et al., 2006], using a statistic called iHS, which measures extended patterns of haplotype homozygosity that are characteristic of selective sweeps. Interestingly, a change in the gene *TYRP1* has also been found to cause a blonde hair phenotype in Melanesians [Kenny et al., 2012]. Another of our top hits is the region containing *SH2B3,* which was identified previously as a candidate for selection in Europe based on both *iHS* and *F_st_* [Pickrell et al., 2009]. This gene contains a nonsynonymous SNP (rs3184504) segregating in Europeans. One of its alleles (the one in the selected haplotype) has been associated with celiac disease and type 1 diabetes [Hunt et al., 2008, Todd et al., 2007] but is also protective against bacterial infection [Zhernakova et al., 2010].

We used Gowinda (v1.12) [Kofler and Schlötterer, 2012] to find enriched Gene Ontology (GO) categories among the regions in the 99.5% highest quantile for each branch score, relative to the rest of the genome (P < 0.05, FDR < 0.3). The significantly enriched categories are listed in Table S4. In the East Asian branch, we find categories related to alcohol catabolism, retinol binding, vitamin metabolism and epidermis development, among others. In the European branch, we find cuticle development and hydrogen peroxide metabolic process as enriched categories. We find no enriched categories in the Eurasian branch that pass the above cutoffs.

### 4.3 Selection in ancestral modern humans

We applied 3P-CLR to modern human data combined with recently sequenced archaic human data. We sought to find selective events that occurred in modern humans after their spit from archaic groups. We used the combined Neanderthal and Denisovan high-coverage genomes [Meyer et al., 2012, Prüfer et al., 2014] as the outgroup population, and, for our two test populations, we used Eurasians (CEU, FIN, GBR, IBS, TSI, CHB, CHS, JPT) and Yoruba (YRI), again from phase 1 of the 1000 Genomes Project Abecasis et al. [2012] (Figure S9.B). As before, we randomly sampled 100 genomes for each of the two daughter populations at each site, and tested for selective events that occurred more anciently than the split of Yoruba and Eurasians, but more recently than the split from Neanderthals. Figure S11 shows an ROC curve for a simulated scenario under these conditions, based on the history of population size changes inferred by PSMC [Li and Durbin, 2011, Prüfer et al., 2014], suggesting we should have power to detect strong (s=0.1) selective events in the ancestral branch of present-day humans. We observe that 3P-CLR(Int) has similar power as XP-CLR and XP-CLR-avg at these time-scales, but is less prone to also detect recent (post-split) events, making it more specific to ancestral sweeps.

We ran 3P-CLR using 0.25 cM windows as above (Figure S13). As before, we selected the top 99.9% windows and merged them together if their corresponding central SNPs were contiguous (Table S5). The top 20 regions are in Table 3. Figure S13 shows that the outliers in the genome-wide distribution of scores are not strong. We wanted to verify that the density of scores was robust to the choice of window size. By using a larger window size (1 cM), we obtained a distribution with slightly more extreme outliers (Figures S12 S13). For that reason, we also show the top hits from this large-window run (Tables S6, 3), using a smaller density of SNPs (200/1cM rather than 100/0.25cM), due to costs in speed. To find putative candidates for the beneficial variants in each region, we queried the catalogs of modern human-specific high-frequency or fixed derived changes that are ancestral in the Neanderthal and/or the Denisova genomes [Castellano et al., 2014, Prüfer et al., 2014] and overlapped them with our regions.

**Table 3.**
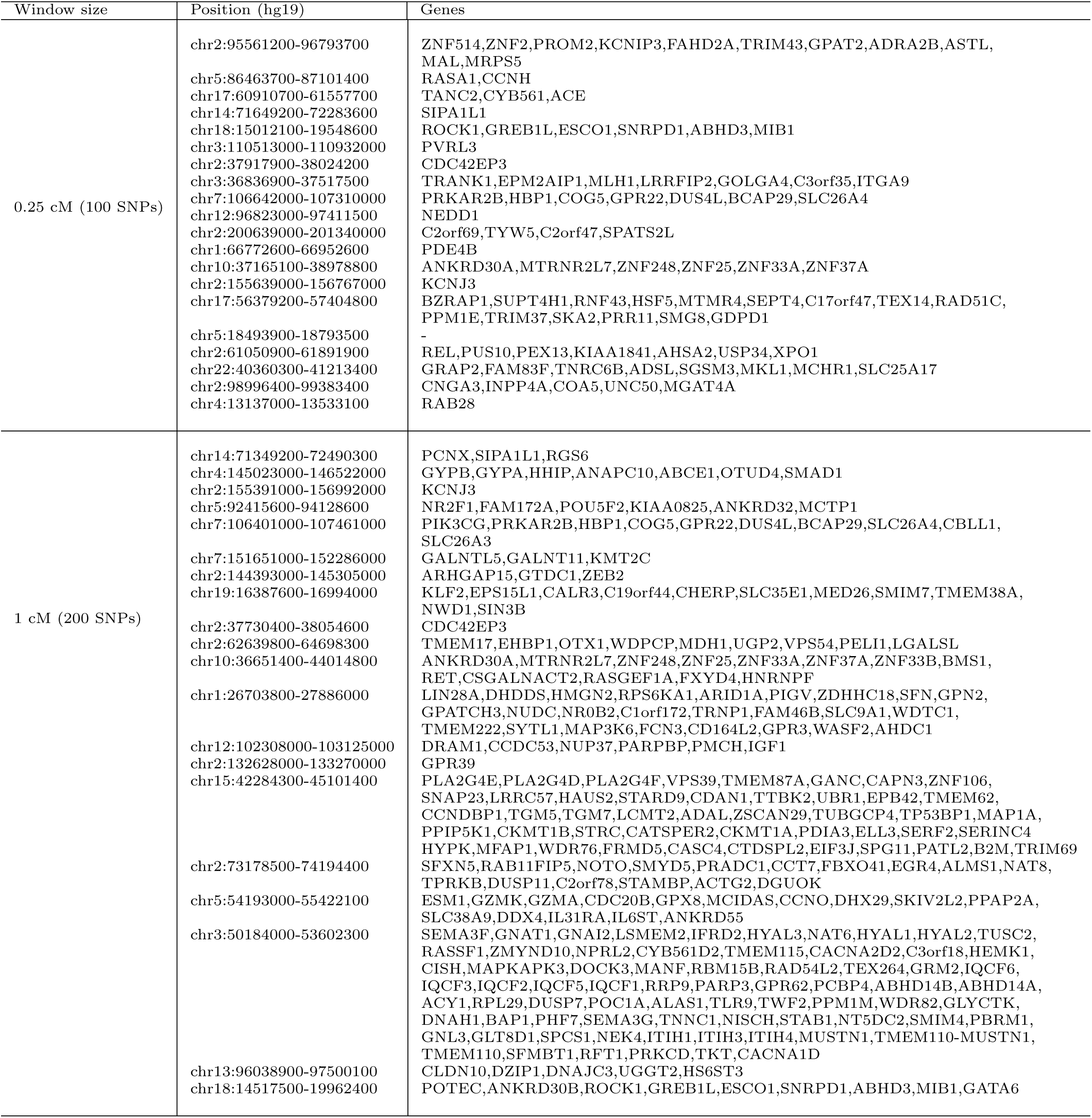
Genes from top 20 candidate regions for the modern human ancestral branch. All positions were rounded to the nearest 100 bp. Windows were merged together if the central SNPs that define them were contiguous.

We found several genes that were identified in previous studies that looked for selection in modern humans after their split from archaic groups [Green et al., 2010, Prüfer et al., 2014], including *SIPA1L1, ANAPC10, ABCE1, RASA1, CCNH, KCNJ3, HBP1, COG5, CADPS2, FAM172A, POU5F2, FGF7, RABGAP1, SMURF1, GABRA2, ALMS1, PVRL3, EHBP1, VPS54, OTX1, UGP2, GTDC1, ZEB2* and *OIT3.* One of our strongest candidate genes among these is *SIPA1L1* (Figure 5.A), which is in the first and the fourth highest-ranking region, when using 1 cM and 0.25 cM windows, respectively. The protein encoded by this gene (E6TP1) is involved in actin cytoskeleton organization and controls neural morphology (UniProt by similarity). Interestingly, it is also a target of degradation of the oncoproteins of high-risk papillomaviruses [Gao et al., 1999].

**Figure 5.**
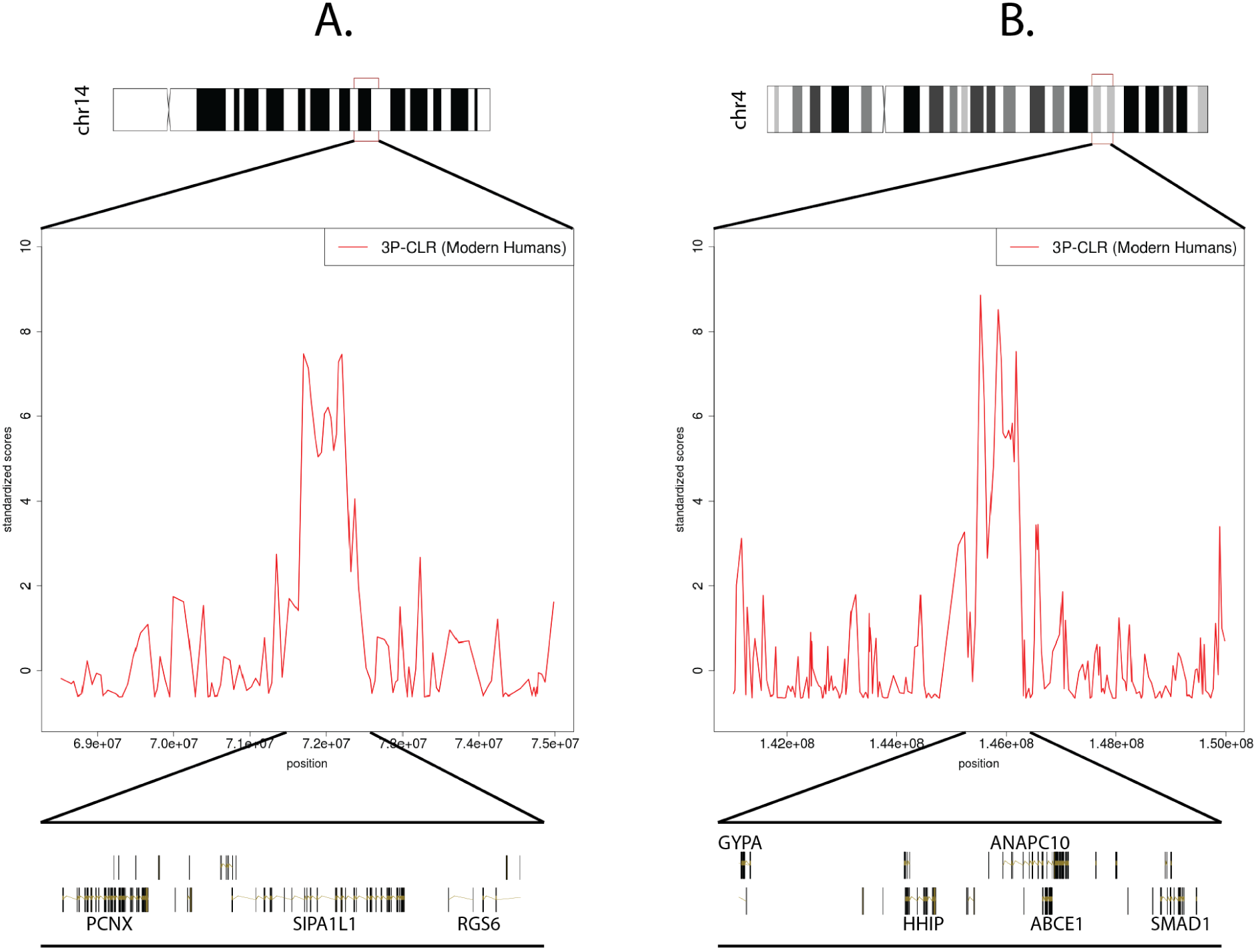
Two of the strongest candidates for selection in the modern human lineage, after the split from Neanderthal and Denisova. We show scores from the 1 cM scan, but the signals persist in the 0.25 cM scan. To make a fair comparison, all 3P-CLR scores were standardized by substracting the chromosome-wide mean from each window and dividing the resulting score by the chromosome-wide standard deviation. A) The region containing *SIPA1L1.* B) The region containing *ANAPC10.* The image was built using the GenomeGraphs package in Bioconductor.

Another candidate gene is *ANAPC10* (Figure 5.B). This gene codes for a core subunit of the cyclosome, which is involved in progression through the cell cycle [Pravtcheva and Wise, 2001], and may play a role in oocyte maturation and human T-lymphotropic virus infection (KEGG pathway [Kanehisa and Goto, 2000]). *ANAPC10* is noteworthy because it was found to be significantly differentially expressed in humans compared to other great apes and macaques: it is up-regulated in the testes [Brawand et al., 2011]. The gene also contains two intronic changes that are fixed derived in modern humans, ancestral in both Neanderthals and Denisovans and that have evidence for being highly disruptive, based on a composite score that combines conservation and regulatory data (PHRED-scaled C-scores > 11 [Kircher et al., 2014, Prüfer et al., 2014]). The changes, however, appear not to lie in any obvious regulatory region [Dunham et al., 2012, Rosenbloom et al., 2011].

We also find *ADSL* among the list of candidates. This gene is known to contain a nonsynonymous change that is fixed in all present-day humans but homozygous ancestral in the Neanderthal genome, the Denisova genome and two Neanderthal exomes [Castellano et al., 2014] (Figure 6.A). It was previously identified as lying in a region with strong support for positive selection in modern humans, using summary statistics implemented in an ABC method [Racimo et al., 2014]. The gene is interesting because it is one of the members of the Human Phenotype ontology category "aggression / hyperactivity" which is enriched for nonsynonymous changes that occurred in the modern human lineage after the split from archaic humans [Castellano et al., 2014, Robinson et al., 2008]. *ADSL* codes for adenylosuccinase, an enzyme involved in purine metabolism [Van Keuren et al., 1987]. A deficiency of adenylosuccinase can lead to apraxia, speech deficits, delays in development and abnormal behavioral features, like hyperactivity and excessive laughter [Gitiaux et al., 2009]. The nonsynonymous mutation (A429V) is in the C-terminal domain of the protein (Figure 6.B) and lies in a highly conserved position (primate PhastCons = 0.953; GERP score = 5.67 [Cooper et al., 2010, Kircher et al., 2014, Siepel et al., 2005]). The ancestral amino acid is conserved across the tetrapod phylogeny, and the mutation is only three residues away from the most common causative SNP for severe adenylosuccinase deficiency [Edery et al., 2003, Kmoch et al., 2000, Maaswinkel-Mooij et al., 1997, Marie et al., 1999, Race et al., 2000]. The change has the highest probability of being disruptive to protein function, out of all the nonsynonymous modern-human-specific changes that lie in the top-scoring regions (C-score = 17.69). While *ADSL* is an interesting candidate and lies in the center of the inferred selected region (Figure 6.A), there are other genes in the region too, including *TNRC6B* and *MKL1. TNRC6B* may be involved in miRNA-guided gene silencing [Meister et al., 2005], while *MKL1* may play a role in smooth muscle differentiation [Du et al., 2004], and has been associated with acute megakaryocytic leukemia [Mercher et al., 2001].

**Figure 6.**
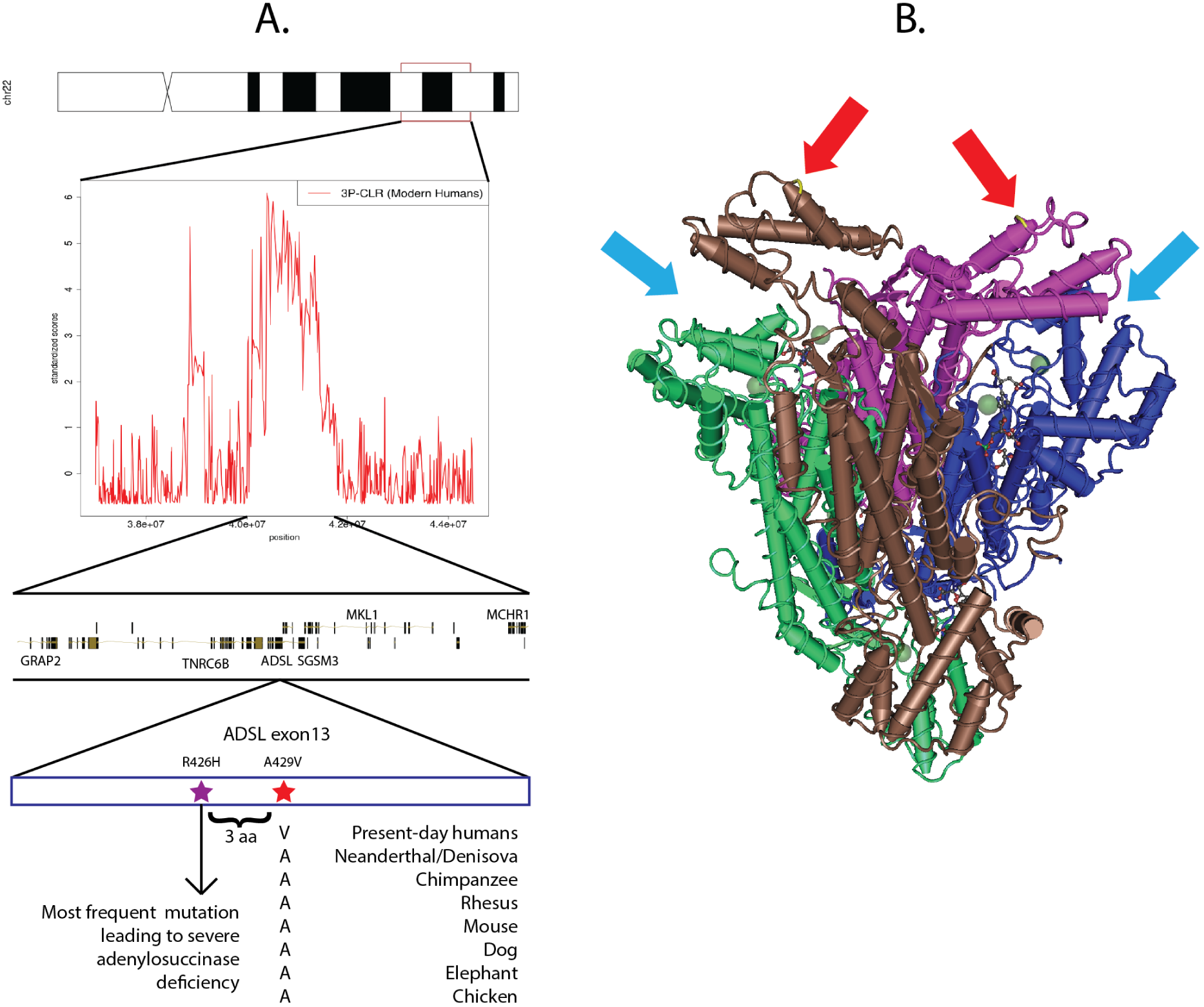
ADSL is a candidate for selection in the modern human lineage, after the split from Neanderthal and Denisova. A) One of the top-scoring regions when running 3P-CLR (0.25 cM windows) on the modern human lineage contains genes *TNRC6B, ADSL, MKL1, MCHR1, SGSM3* and *GRAP2.* The most disruptive nonsynonymous modern-human-specific change in the entire list of top regions is in an exon of *ADSL* and is fixed derived in all present-day humans but ancestral in archaic humans. It is highly conserved across tetrapods and lies only 3 residues away from the most common mutation leading to severe adenylosuccinase deficiency. B) The *ADSL* gene codes for a tetrameric protein. The mutation is in the C-terminal domain of each tetrameric unit (red arrows), which are near the active sites (light blue arrows). Scores in panel A were standardized using the chromosome-wide mean and standard deviation. Vertebrate alignments were obtained from the UCSC genome browser (Vertebrate Multiz Alignment and Conservation track) and the image was built using the GenomeGraphs package in Bioconductor and Cn3D.

*RASA1* was also a top hit in a previous scan for selection [Green et al., 2010], and was additionally inferred to have evidence in favor of selection in Racimo et al. [2014]. The gene codes for a protein involved in the control of cellular differentiation [Trahey et al., 1988], and has a modern human-specific fixed nonsynonymous change (G70E). Human diseases associated with *RASA1* include basal cell carcinoma [Friedman et al., 1993] and arteriovenous malformation [Eerola et al., 2003, Hershkovitz et al., 2008].

The *GABA_A_* gene cluster in chromosome 4p12 is also among the top regions. The gene within the putatively selected region codes for a subunit *(GABRA2*) of the *GABA_A_* receptor, which is a ligand-gated ion channel that plays a key role in synaptic inhibtion in the central nervous system (see review by Whiting et al. [1999]). *GABRA2* is significantly associated with risk of alcohol dependence in humans [Edenberg et al., 2004], perception of pain [Knabl et al., 2008] and asthma [Xiang et al., 2007].

Two other candidate genes that may be involved in brain development are *FOXG1* and *CADPS2. FOXG1* was not identified in any of the previous selection scans, and codes for a protein called forkhead box G1, which plays an important role during brain development. Mutations in this gene are associated with a slow-down in brain growth during childhood resulting in microcephaly, which in turn causes various intellectual disabilities [Ariani et al., 2008, Mencarelli et al., 2010]. *CADPS2* was identified in Green et al. [2010] as a candidate for selection, and has been associated with autism [Sadakata and Furuichi, 2010]. The gene has been suggested to be specifically important in the evolution of all modern humans, as it was not found to be selected earlier in great apes or later in particular modern human populations [Crisci et al., 2011].

Finally, we find a signal of selection in a region containing the gene *EHBP1* and *OTX1.* This region was identified in both of the two previous scans for modern human selection [Green et al., 2010, Prüfer et al., 2014]. *EHBP1* codes for a protein involved in endocytic trafficking [Guilherme et al., 2004] and has been associated with prostate cancer [Gudmundsson et al., 2008]. *OTX1* is a homeobox family gene that may play a role in brain development [Gong et al., 2003]. Interestingly, *EHBP1* contains a single-nucleotide intronic change (chr2:63206488) that is almost fixed in all present-day humans and homozygous ancestral in Neanderthal and Denisova [Prüfer et al., 2014]. This change is also predicted to be highly disruptive (C-score = 13.1) and lies in a position that is extremely conserved across primates (PhastCons = 0.942), mammals (PhastCons = 1) and vertebrates (PhastCons = 1). The change is 18 bp away from the nearest splice site and overlaps a VISTA conserved enhancer region (element 1874) [Pennacchio et al., 2006], suggesting a putative regulatory role for the change.

We again used Gowinda [Kofler and Schlötterer, 2012] to find enriched GO categories among the regions with high 3P-CLR scores in the Modern Human branch. The significantly enriched categories (P < 0.05, FDR < 0.3) are listed in Table S4. We find several GO terms related to the regulation of the cell cycle, T cell migration and intracellular transport.

We overlapped the genome-wide association studies (GWAS) database [Li et al., 2011, Welter et al., 2014] with the list of fixed or high-frequency modern human-specific changes that are ancestral in archaic humans [Prüfer et al., 2014] and that are located within our top putatively selected regions in modern humans (Tables S7 and S8 for the 0.25 cM and 1 cM scans, respectively). None of the resulting SNPs are completely fixed derived, because GWAS can only yield associations from sites that are segregating. We find several SNPs in the *RAB28* gene [Dunham et al., 2012, Rosenbloom et al., 2011], which are significantly associated with obesity [Paternoster et al., 2011]. We also find a SNP with a high C-score (rs10171434) associated with urinary metabolites [Suhre et al., 2011] and suicidal behavior in patients with mood disorders [Perlis et al., 2010]. The SNP is located in an enhancer regulatory freature [Dunham et al., 2012, Rosenbloom et al., 2011] located between genes *PELI1* and *VPS54,* in the same putatively selected region as genes *EHBP1* and *OTX1* (see above). Finally, there is a highly C-scoring SNP (rs731108) that is associated with renal cell carcinoma [Henrion et al., 2013]. This SNP is also located in an enhancer regulatory feature [Dunham et al., 2012, Rosenbloom et al., 2011], in an intron of *ZEB2.* In this last case, though, only the Neanderthal genome has the ancestral state, while the Denisova genome carries the modern human variant.

## 5 Discussion

We have developed a new method called 3P-CLR, which allows us to detect positive selection along the genome. The method is based on an earlier test (XP-CLR [Chen et al., 2010]) that uses linked allele frequency differences between two populations to detect population-specific selection. However, unlike XP-CLR, 3P-CLR can allow us to distinguish between selective events that occurred before and after the split of two populations. Our method has some similiarities to an earlier method developed by Schlebusch et al. [2012], which used an *F_st_*-like score to detect selection ancestral to two populations. In that case, though, the authors used summary statistics and did not explicitly model the process leading to allele frequency differentiation. It is also similar to a more recent method [Fariello et al., 2013] that models differences in haplotype frequencies between populations, while accounting for population structure.

We used our method to confirm previously found candidate genes in particular human populations, like *EDAR, TYRP1* and *CYP26B1,* and find some novel candidates too (Tables S1, S2, S3). Additionally, we can infer that certain genes, which were previously known to have been under selection in East Asians (like *SPAG6*), are more likely to have undergone a sweep in the population ancestral to both Europeans and East Asians than in East Asians only. We find that genes involved in epidermis development and alcohol catabolism are particularly enriched among the East Asian candidate regions, while genes involved in peroxide catabolism and cuticle development are enriched in the European branch. This suggests these biological functions may have been subject to positive selection in recent times.

We also used 3P-CLR to detect selective events that occurred in the ancestors of modern humans, after their split from Neanderthals and Denisovans (Table S5). These events could perhaps have led to the spread of phenotypes that set modern humans apart from other hominin groups. We find several intersting candidates, like *SIPA1L1, ADSL, RASA1, OTX1, EHBP1, FOXG1, RAB28* and *ANAPC10,* some of which were previously detected using other types of methods [Green et al., 2010, Prüfer et al., 2014, Racimo et al., 2014]. We also find an enrichment for GO categories related to cell cycle regulation and T cell migration among the candidate regions, suggesting that these biological processes might have been affected by positive selection after the split from archaic humans.

An advantage of differentiation-based tests like XP-CLR and 3P-CLR is that, unlike other patterns detected by tests of neutrality (like extended haplotype homozygostiy, [Sabeti et al., 2002]) that are exclusive to hard sweeps, the patterns that both XP-CLR and 3P-CLR are tailored to find are based on regional allele frequency differences between populations. These patterns can also be produced by soft sweeps from standing variation or by partial sweeps [Chen et al., 2010], and there is some evidence that the latter phenomena may have been more important than classic sweeps during human evolutionary history [Hernandez et al., 2011].

Another advantage of both XP-CLR and 3P-CLR is that they do not rely on an arbitrary division of genomic space. Unlike other methods which require the partition of the genome into small windows of fixed size, our composite likelihood ratios can theoretically be computed over windows that are as big as each chromosome, while only switching the central candidate site at each window. This is because the likelihood ratios use the genetic distance to the central SNP as input. SNPs that are very far away from the central SNP will not contribute much to the likelihood function of both the neutral and the selection models, while those that are close to it will. In the interest of speed, we heuristically limit the window size in our implementation, and use less SNPs when calculating likelihoods over larger windows. Nevertheless, these parameters can be arbitrarily adjusted by the user as needed, and if enough computing resources are available. The use of genetic distance in the likelihood function also allows us to take advantage of the spatial distribution of SNPs as an additional source of information, rather than only relying on patterns of population differentiation restricted to tightly linked SNPs.

3P-CLR also has an advantage over HMM-based selection methods, like the one implemented in Prüfer et al. [2014]. The likelihood ratio scores obtained from 3P-CLR can provide an idea of how credible a selection model is for a particular region, relative to the rest of the genome. The HMM-based method previously used to scan for selection in modern humans [Prüfer et al., 2014] can only rank putatively selected regions by genetic distance, but cannot output a statistical measure that may indicate how likely each region is to have been under selection in ancient times. In contrast, 3P-CLR provides a composite likelihood ratio score, which allows for a statistically rigorous way to compare the neutral model and a specific selection model (for example, recent or ancient selection).

The outliers from Figure S10 have much higher scores (relative to the rest of the genome) than the outliers from Figure S13. This may be due to both the difference in time scales in the two sets of tests and to the uncertainty that comes from estimating outgroup allele frequencies using only two archaic genomes. This pattern can also be observed in Figure S14, where the densities of the scores looking for patterns of ancient selection (3P-CLR Modern Human and 3P-CLR Eurasia) have much shorter tails than the densities of scores looking for patterns of recent selection (3P-CLR Europe and 3P-CLR East Asia). Simulations show that 3P-CLR(Int) score distributions are naturally shorter than 3P-CLR(A) scores (Figure S15), which could explain the short tail of the 3P-CLR Eurasia distribution. Additionally, the even shorter tail in the distribution of 3P-CLR Modern Human scores may be a consequence of the fact that the split times of the demographic history in that case are older than the split times in the Eurasian tree, as simulations show that ancient split times tend to further shorten the tail of the 3P-CLR score distribution (Figure S15). We note, though, that using a larger window size produces a larger number of strong outliers (Figure S12).

A limitation of composite likelihood ratio tests is that the composite likelihood calculated for each model under comparison is obtained from a product of individual likelihoods at each site, and so it underestimates the correlation that exists between SNPs due to linkage effects [Chen et al., 2010, Lindsay, 1988, Pace et al., 2011, Varin et al., 2011]. One way to partially mitigate this problem is by using corrective weights based on linkage disequilibrium (LD) statistics calculated on the outgroup population [Chen et al., 2010]. Our implementation of 3P-CLR allows the user to incorporate such weights, if appropriate LD statistics are available from the outgroup. However, in cases where these are unreliable, it may not be possible to correct for this (for example, when only a few unphased genomes are available, as in the case of the Neanderthal and Denisova genomes).

While 3P-CLR relies on integrating over the possible allele frequencies in the ancestors of populations *a* and *b* (formula 10), one could envision using ancient DNA to avoid this step. Thus, if enough genomes could be sampled from that ancestral population that existed in the past, one could use the sample frequency in the ancient set of genomes as a proxy for the ancestral population frequency. This may soon be possible, as several early modern human genomes have already been sequenced in recent years [Fu et al., 2014, Lazaridis et al., 2014, Seguin-Orlando et al., 2014].

Though we have focused on a three-population model in this manuscript, it should be straightforward to expand our method to a larger number of populations, albeit with additional costs in terms of speed and memory. 3P-CLR relies on a similar framework to the demographic inference method implemented in TreeMix [Pickrell and Pritchard, 2012], which can estimate population trees that include migration events, using genome-wide data. With a more complex modeling framework, it may be possible to estimate the time and strength of selective events with better resolution and using more populations, and also to incorporate additional demographic forces, like continuous migration between populations or pulses of admixture.

## Acknowledgments

We thank Montgomery Slatkin, Rasmus Nielsen, Joshua Schraiber, Nicolas Duforet-Frebourg, Emilia Huerta-Sánchez, Hua Chen, Benjamin Peter, Nick Patterson, David Reich, Joachim Hermisson, Graham Coop and members of the Slatkin and Nielsen labs for helpful advice and discussions. We also thank two anonymous reviewers for their helpful comments. This work was supported by NIH grant R01-GM40282 to Montgomery Slatkin.

## Supplementary Tables

**Table S1.**
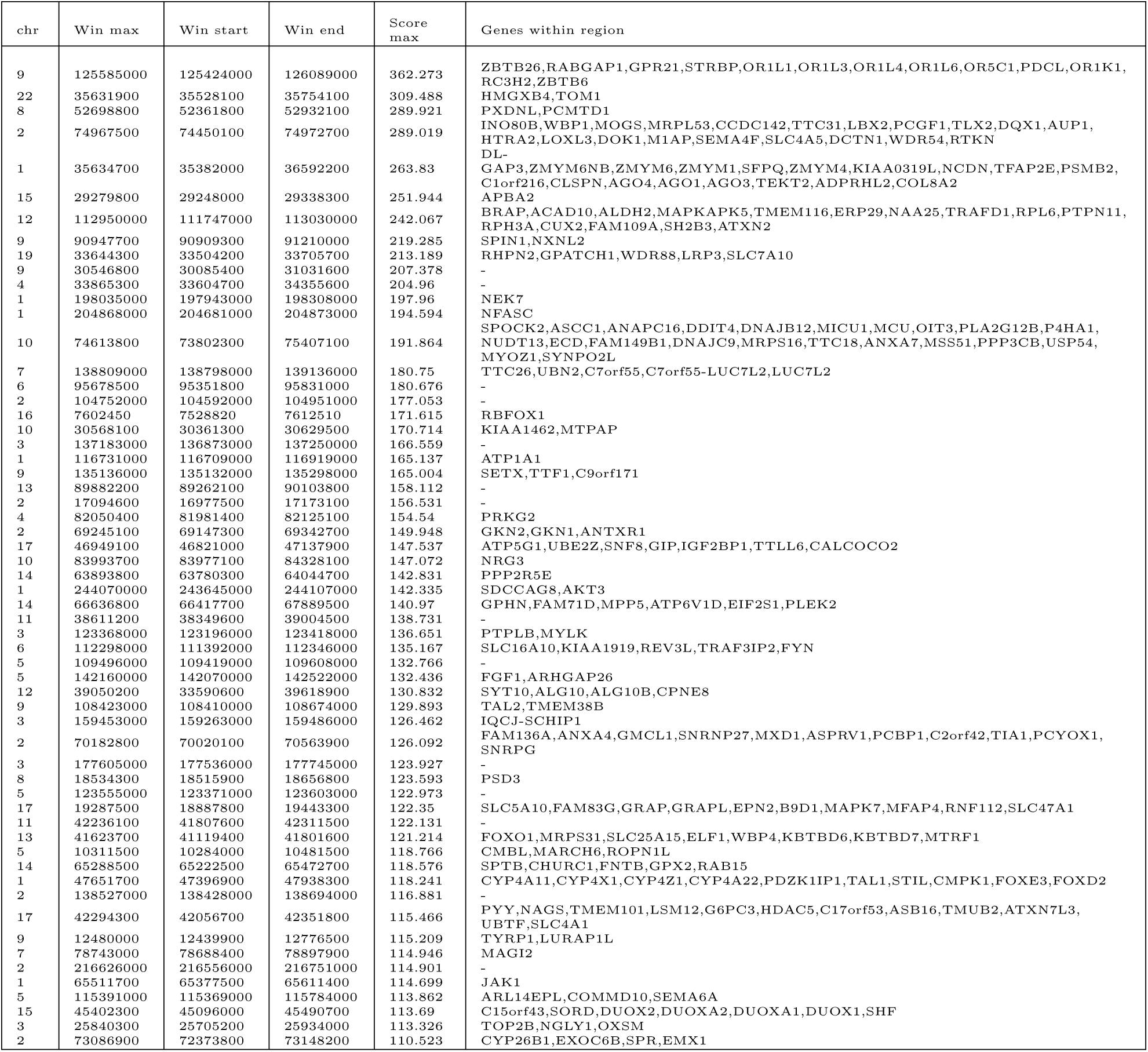
Top hits for 3P-CLR run on the European terminal branch, using Yoruba as the outgroup. We show the windows in the top 99.9% quantile of scores. Windows were merged together if the central SNPs that define them were contiguous. Win max = Location of window with maximum score. Win start = left-most end of left-most window for each region. Win end = right-most end of right-most window for each region. All positions were rounded to the nearest 100 bp. Score max = maximum score within region.

**Table S2.**
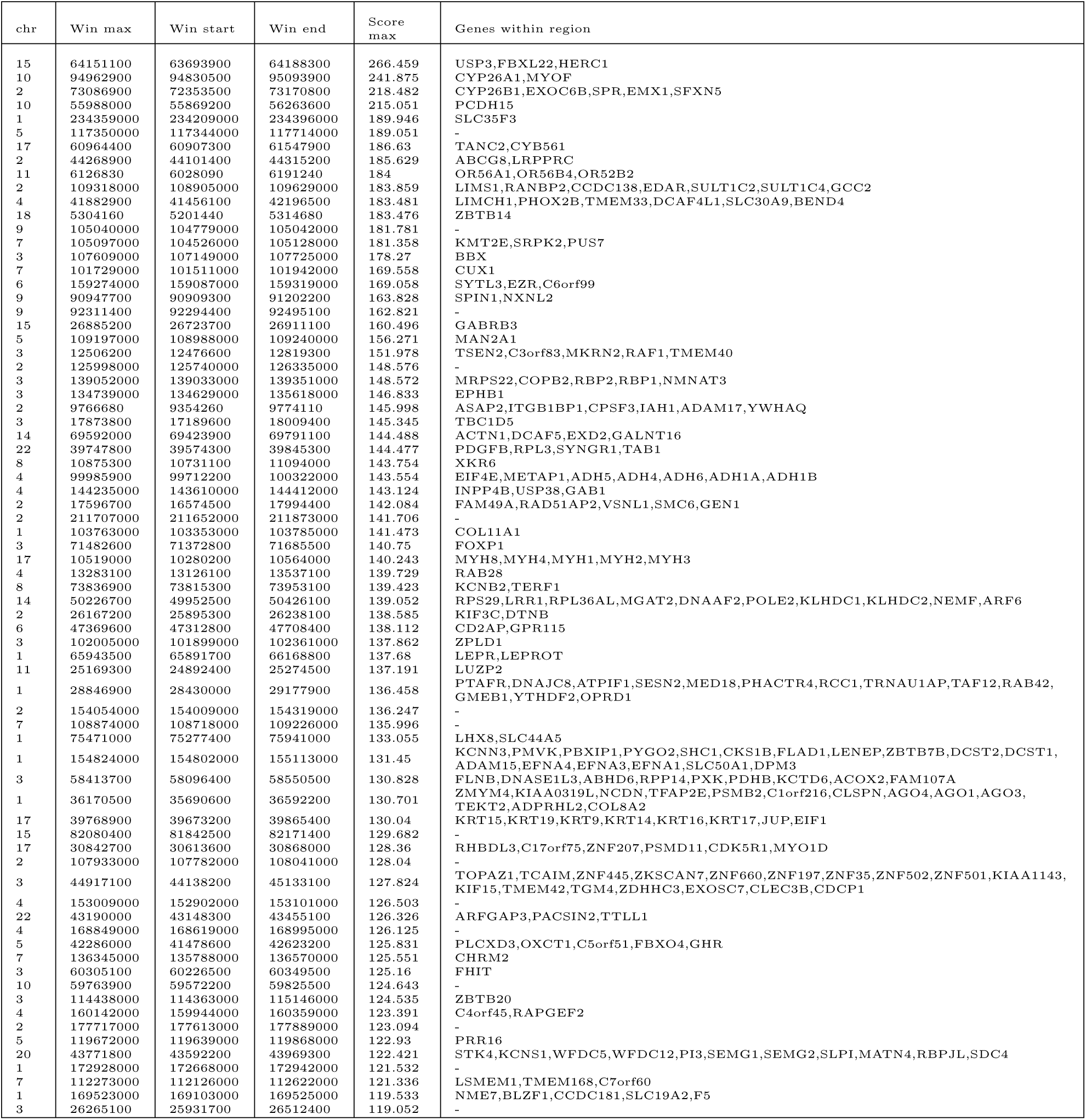
Top hits for 3P-CLR run on the East Asian terminal branch, using Yoruba as the outgroup. We show the windows in the top 99.9% quantile of scores. Windows were merged together if the central SNPs that define them were contiguous. Win max = Location of window with maximum score. Win start = left-most end of left-most window for each region. Win end = right-most end of right-most window for each region. All positions were rounded to the nearest 100 bp. Score max = maximum score within region.

**Table S3.**
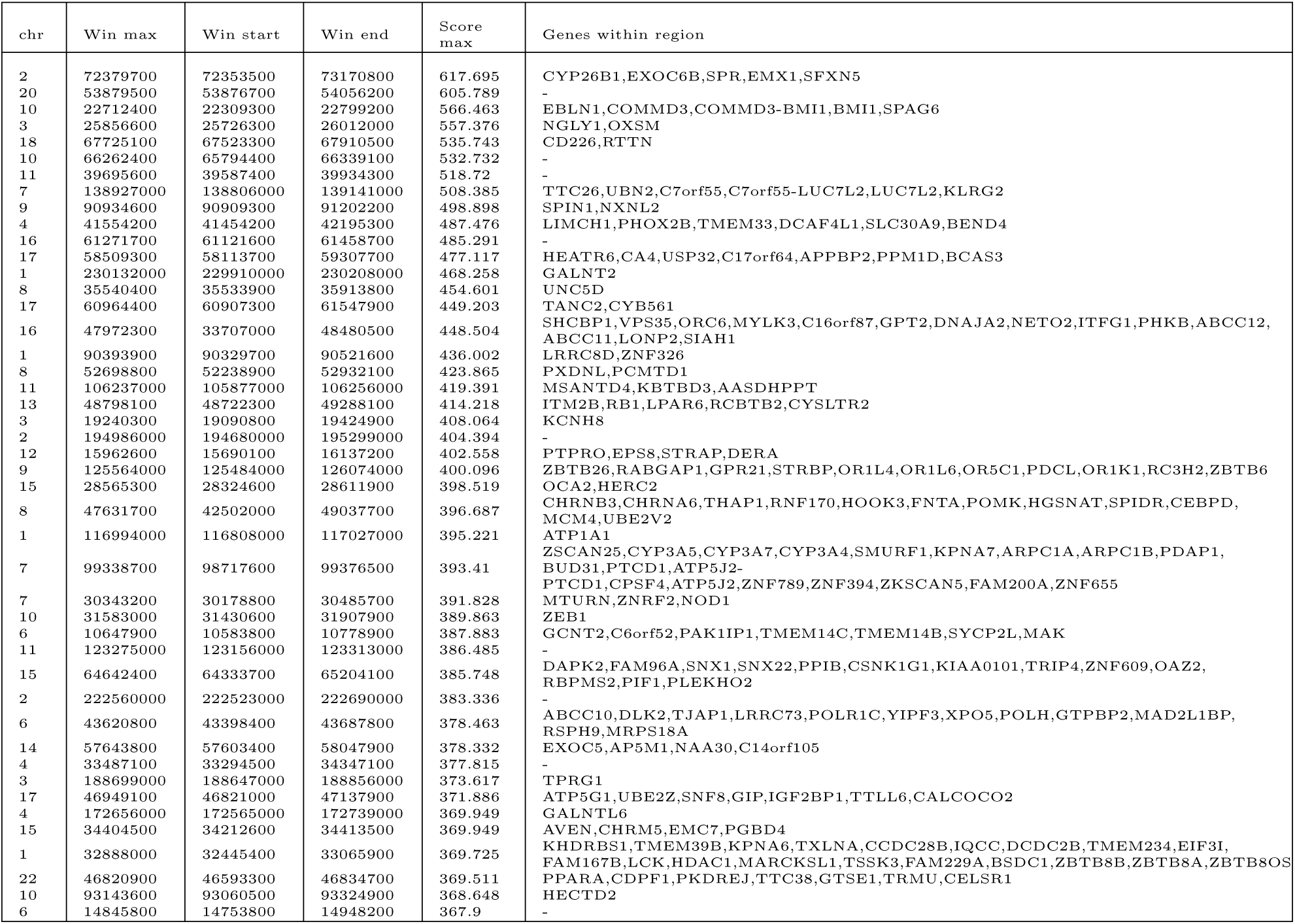
Top hits for 3P-CLR run on the Eurasian ancestral branch, using Yoruba as the outgroup. We show the windows in the top 99.9% quantile of scores. Windows were merged together if the central SNPs that define them were contiguous. Win max = Location of window with maximum score. Win start = left-most end of left-most window for each region. Win end = right-most end of right-most window for each region. All positions were rounded to the nearest 100 bp. Score max = maximum score within region.

**Table S4.**
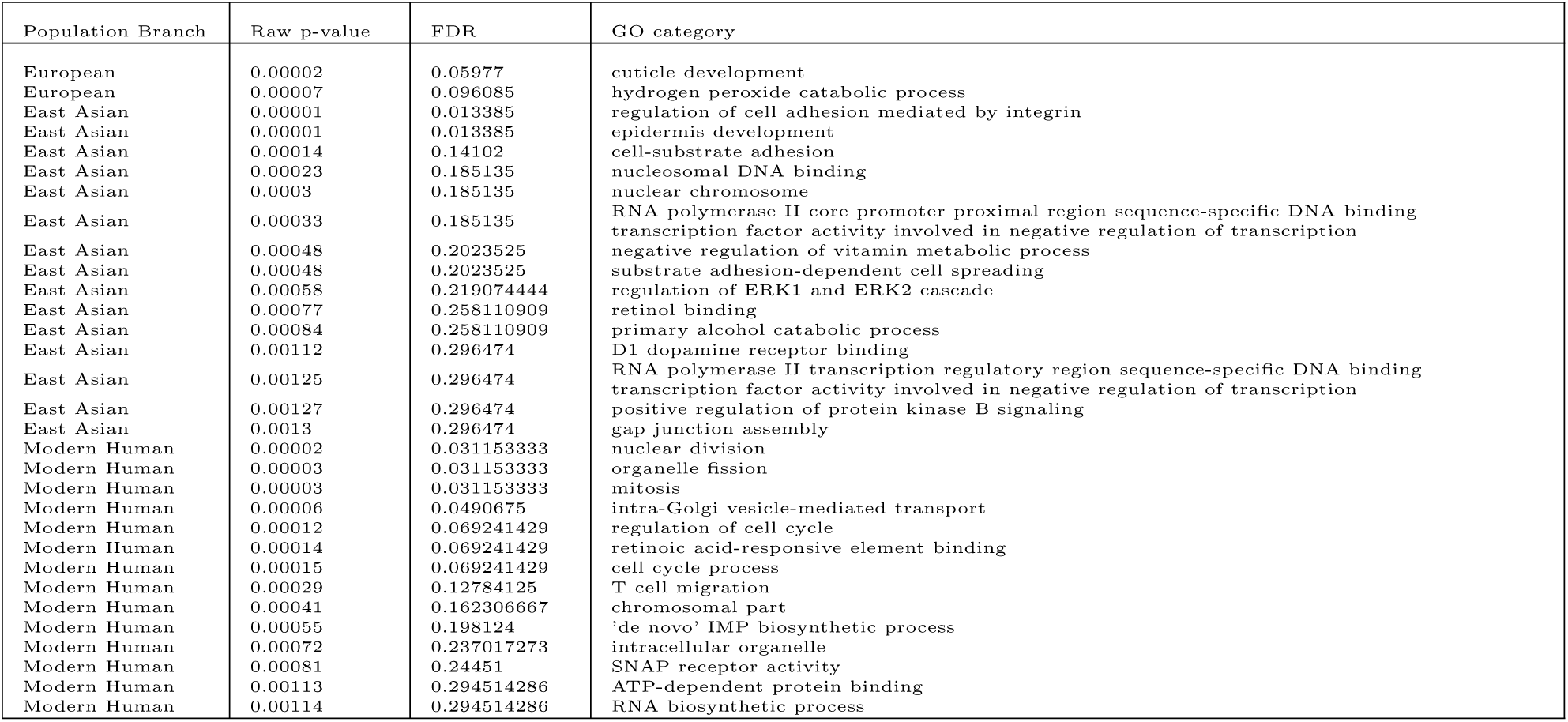
Enriched GO categories in the European, East Asian and Modern Human branches. We tested for ontology enrichment among the regions in the 99.5% quantile of the 3P-CLR scores for each population branch (P < 0.05, FDR < 0.3). The Eurasian branch did not have any category that passed these cutoffs.

**Table S5.**
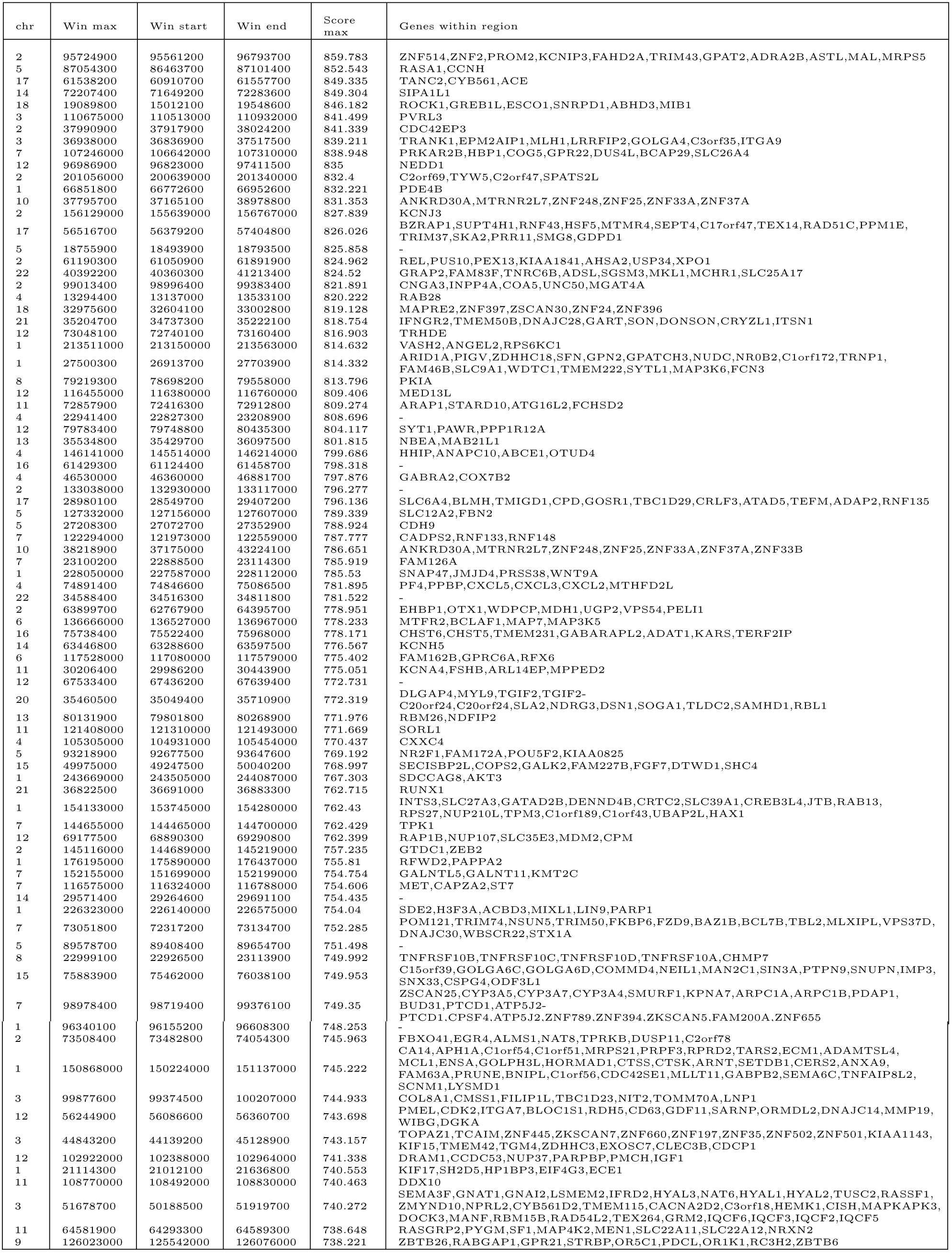
Top hits for 3P-CLR run on the ancestral branch to Eurasians and Yoruba, using archaic humans as the outgroup and 0.25 cM windows. We show the windows in the top 99.9% quantile of scores. Windows were merged together if the central SNPs that define them were contiguous. Win max = Location of window with maximum score. Win start = left-most end of left-most window for each region. Win end = right-most end of right-most window for each region. All positions were rounded to the nearest 100 bp. Score max = maximum score within region.

**Table S6.**
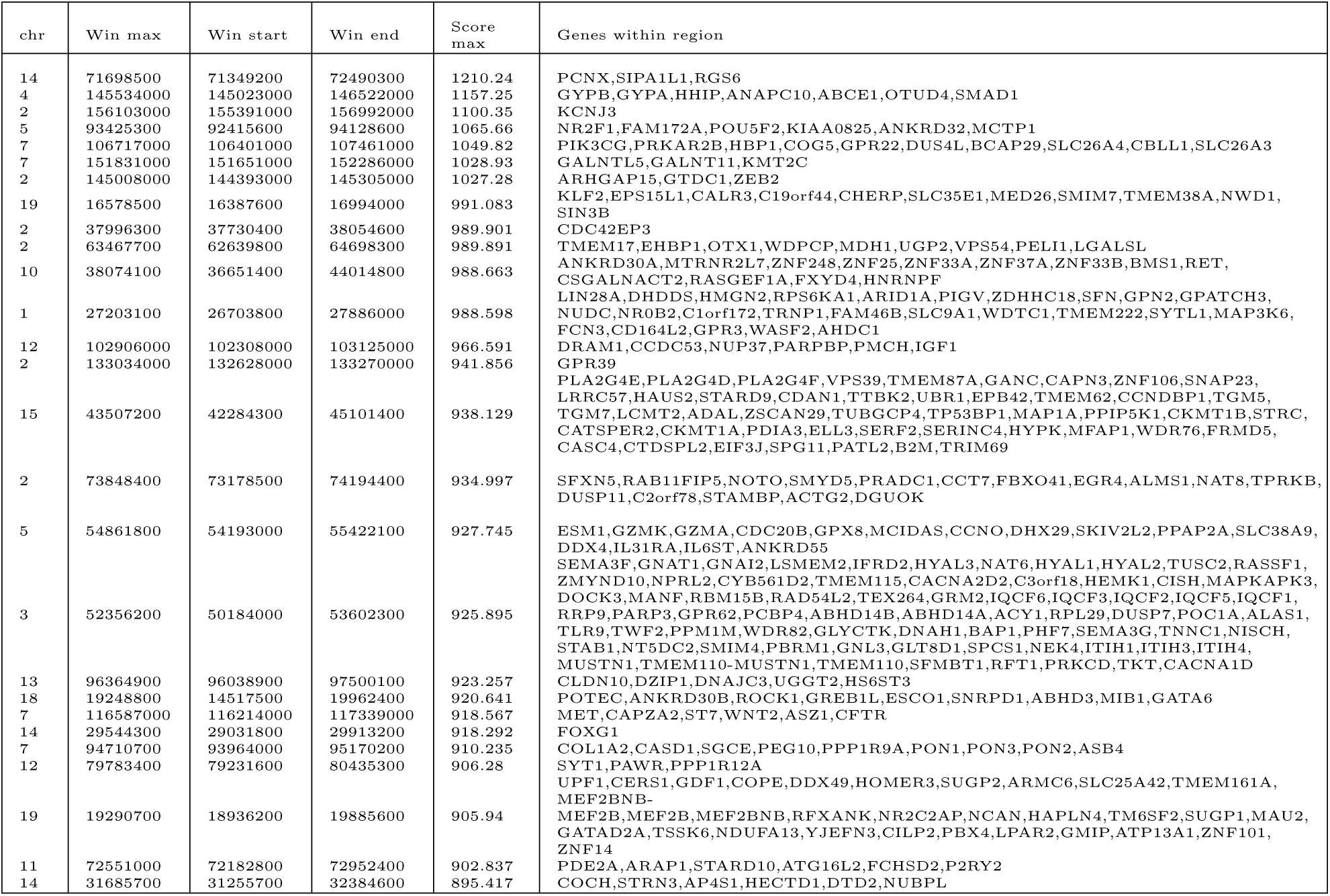
Top hits for 3P-CLR run on the ancestral branch to Eurasians and Yoruba, using archaic humans as the outgroup and 1 cM windows. We show the windows in the top 99.9% quantile of scores. Windows were merged together if the central SNPs that define them were contiguous. Win max = Location of window with maximum score. Win start = left-most end of left-most window for each region. Win end = right-most end of right-most window for each region. All positions were rounded to the nearest 100 bp. Score max = maximum score within region.

**Table S7.**
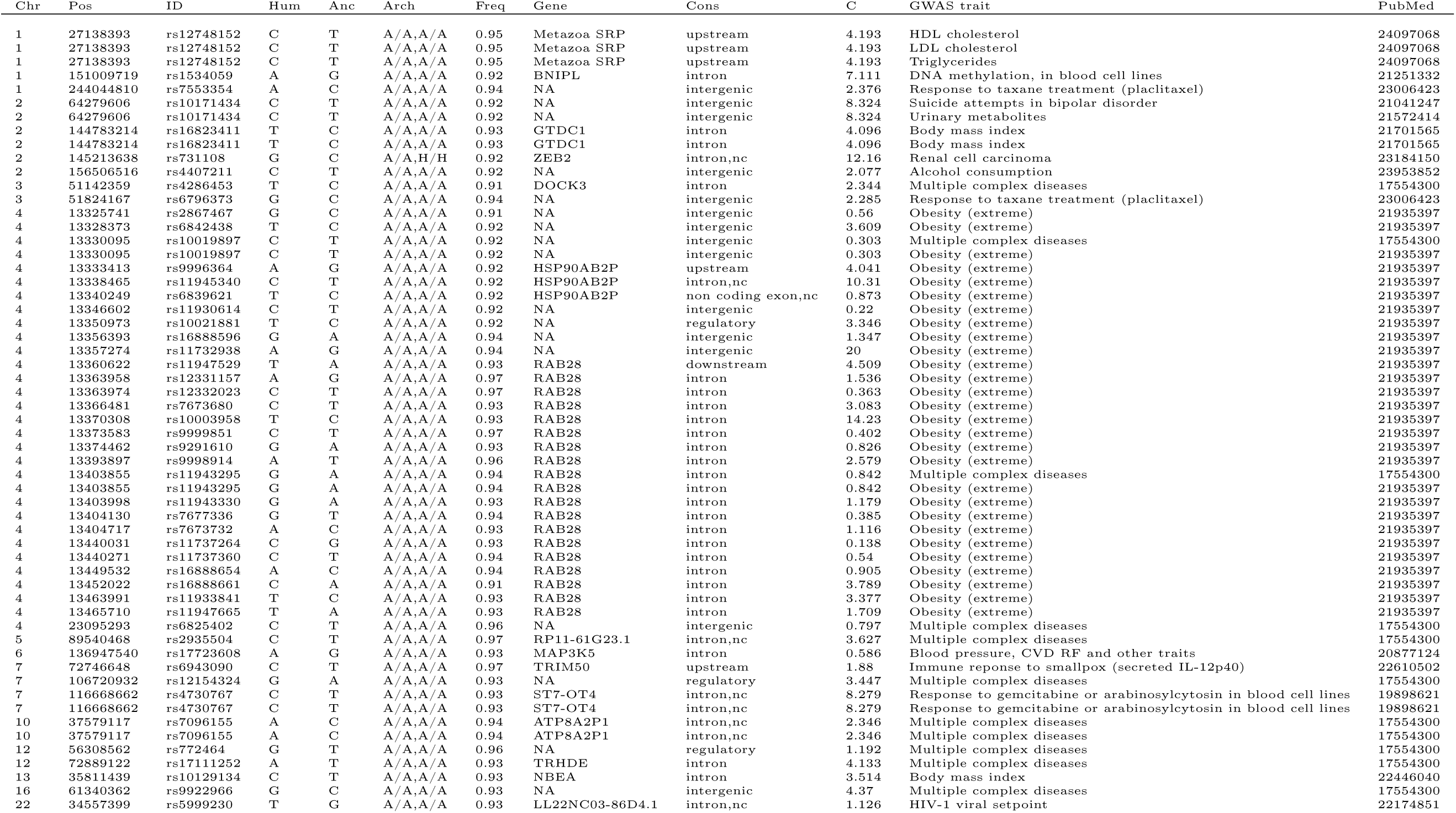
Overlap between GWAS catalog and catalog of modern human-specific high-frequency changes in the top modern human selected regions (0.25 cM scan). Chr = chromosome. Pos = position (hg19). ID = SNP rs ID. Hum = Present-day human major allele. Anc = Human-Chimpanzee ancestor allele. Arch = Archaic human allele states (Altai Neanderthal, Denisova) where H=human-like allele and A=ancestral allele. Freq = present-day human derived frequency. Cons = consequence. C = C-score. PubMed = PubMed article ID for GWAS study.

**Table S8.**
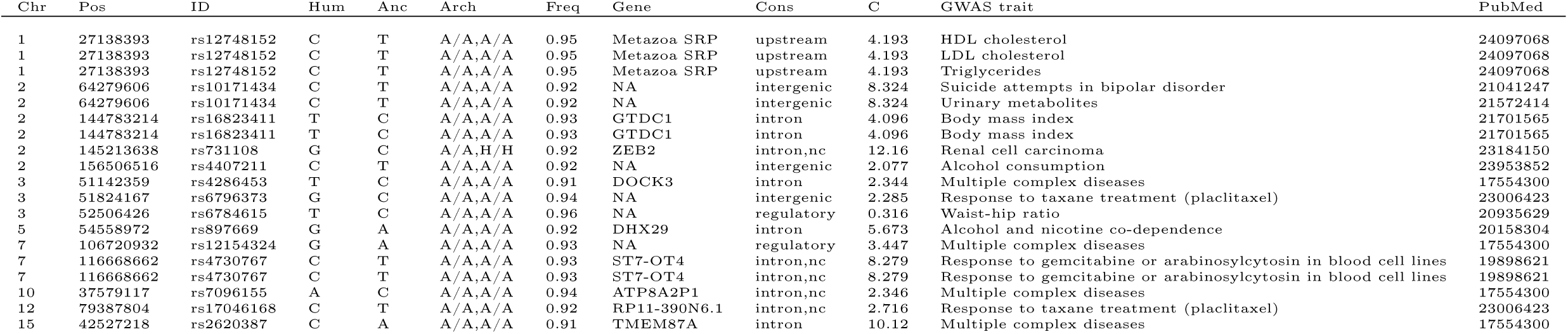
Overlap between GWAS catalog and catalog of modern human-specific high-frequency changes in the top modern human selected regions (1 cM scan). Chr = chromosome. Pos = position (hg19). ID = SNP rs ID. Hum = Present-day human major allele. Anc = Human-Chimpanzee ancestor allele. Arch = Archaic human allele states (Altai Neanderthal, Denisova) where H=human-like allele and A=ancestral allele. Freq = present-day human derived frequency. Cons = consequence. C = C-score. PubMed = PubMed article ID for GWAS study.

## Supplementary Figures

**Figure S1.**
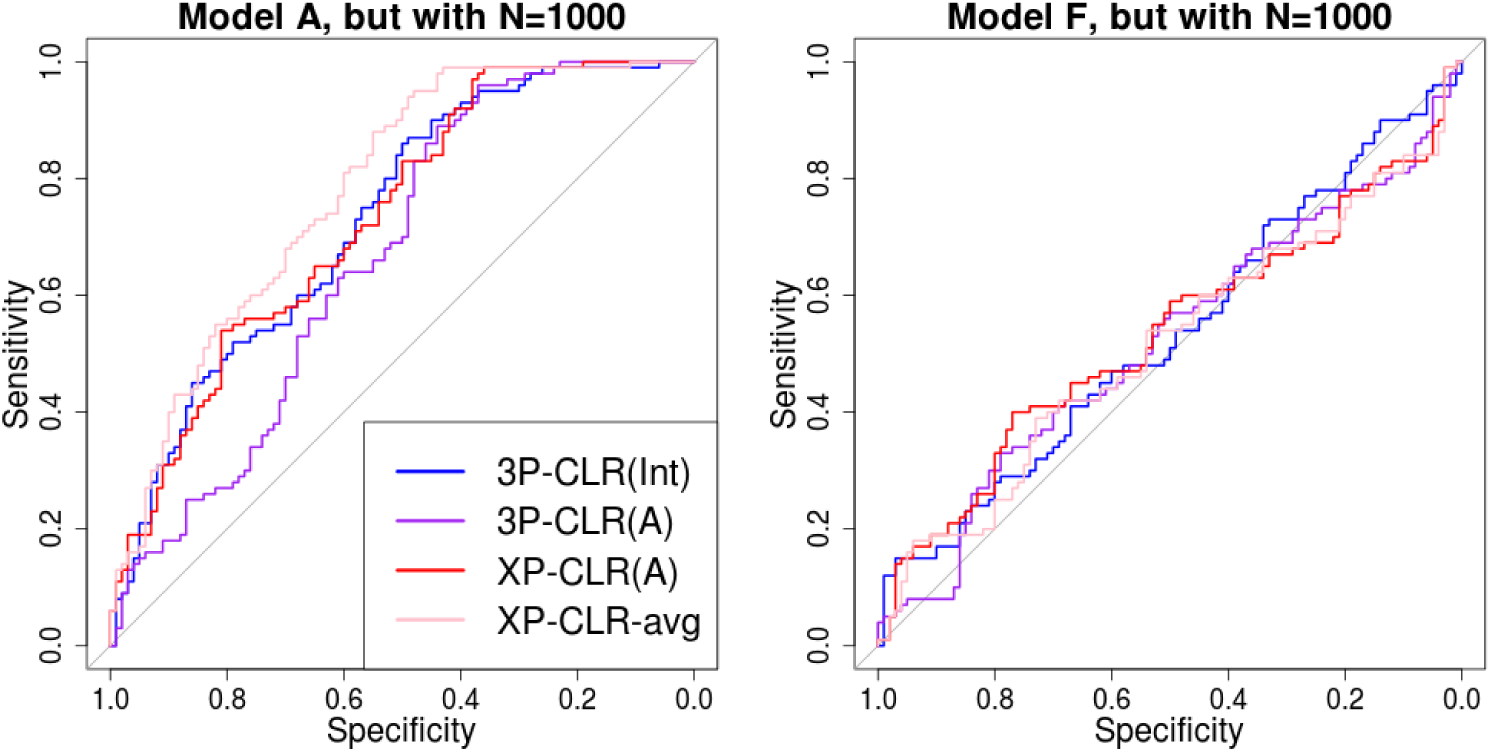
ROC curves for performance of 3P-CLR(Int), 3P-CLR(A) and two variants of XP-CLR in detecting selective sweeps that occurred before the split of two populations *a* and *b*, under two demographic models where the population size is extremely small (*N_e_* = 1,000).

**Figure S2.**
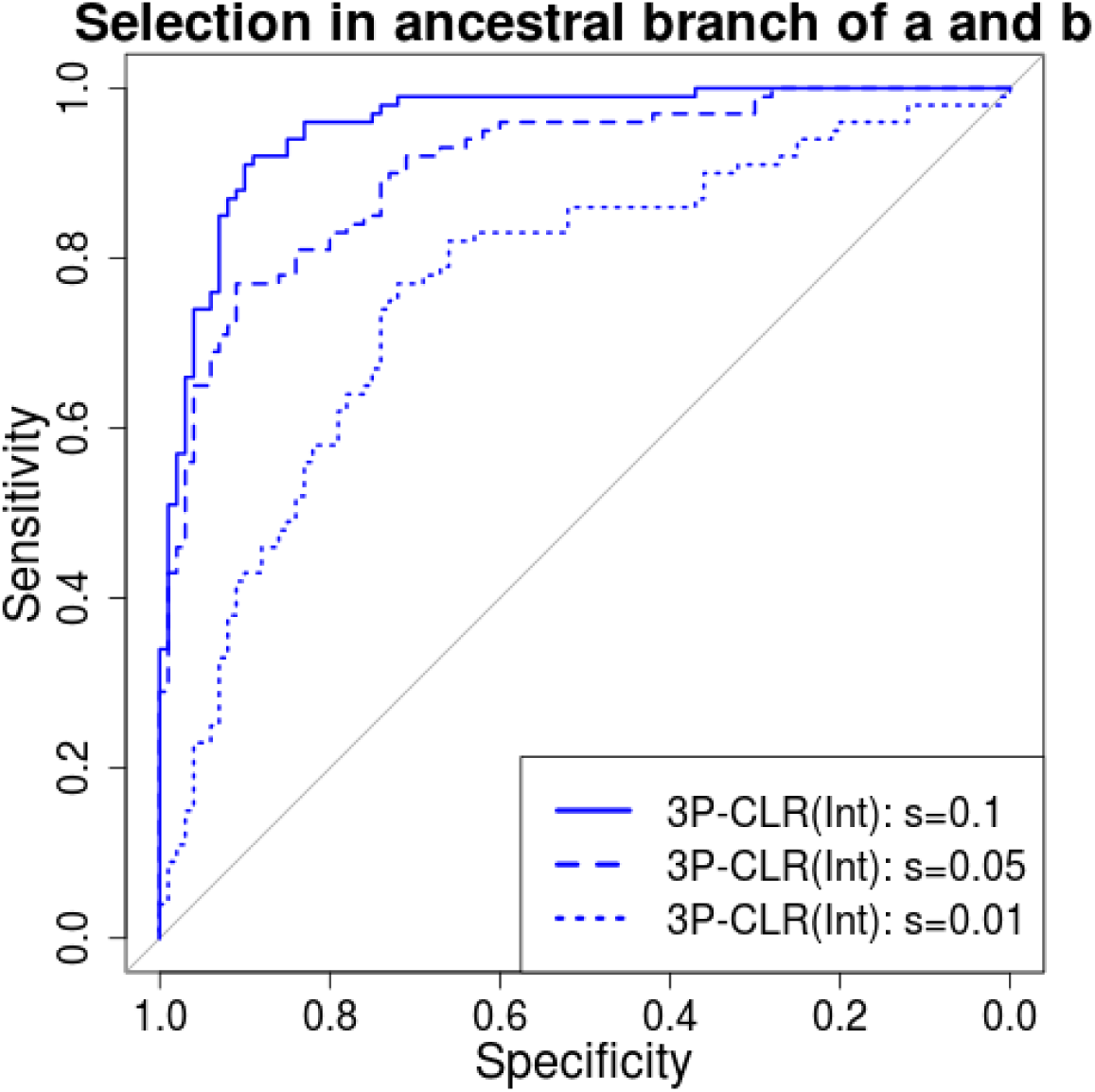
Performance of 3P-CLR(Int) for a range of selection coefficients. We used the demographic history from model B (Table 1) but extended the most ancient split time by 4,000 generations. The reason for this is that we wanted the internal branch to be long enough for it to be easy to sample simulations in which the beneficial allele fixed before the split of populations *a* and *b*, even for weak selection coefficients.

**Figure S3.**
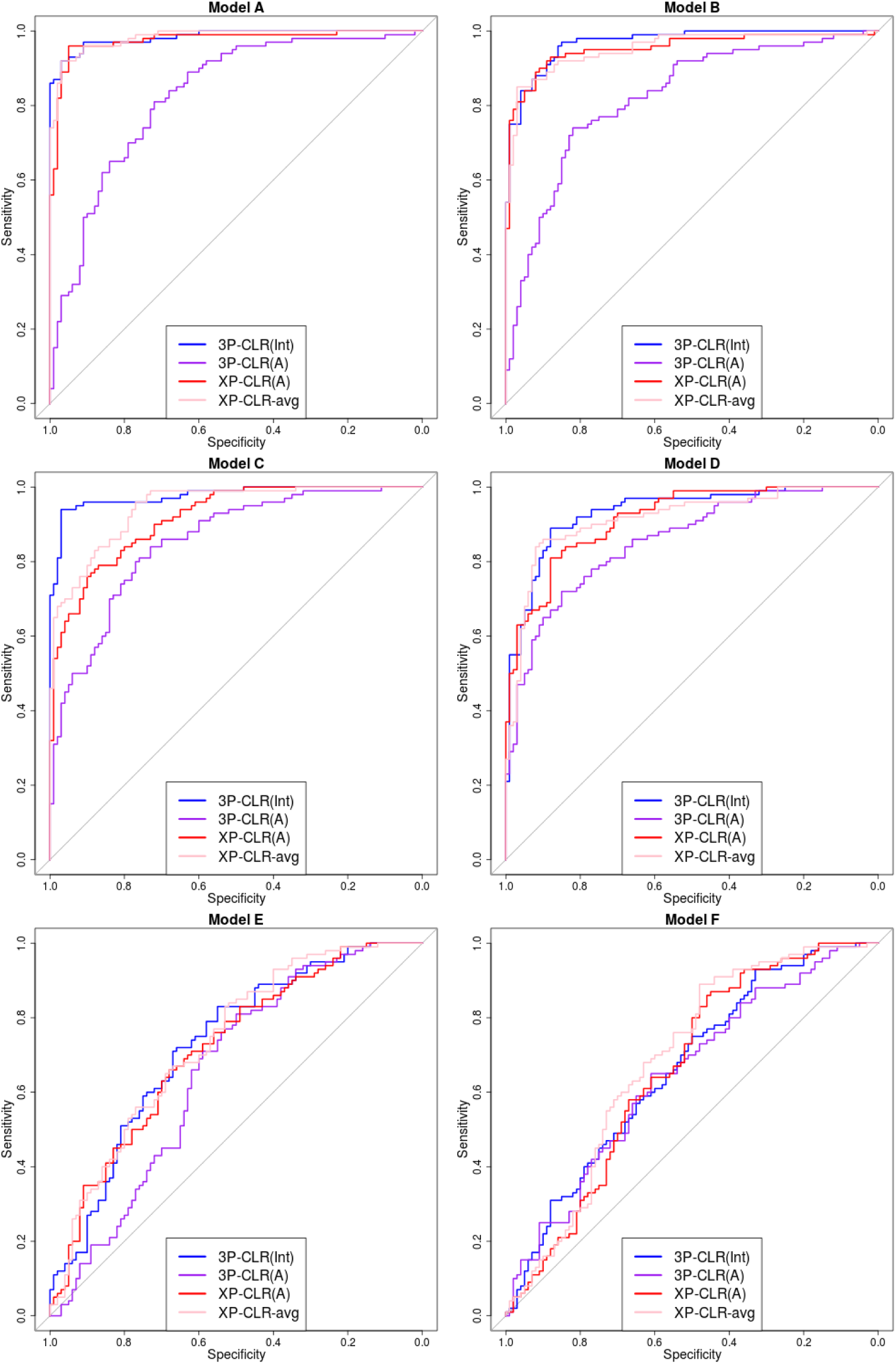
ROC curves for performance of 3P-CLR(Int), 3P-CLR(A) and two variants of XP-CLR in detecting selective sweeps that occurred before the split of two populations *a* and *b*, under different demographic models. In this case, the outgroup panel from population *c* contained 10 haploid genomes. The two sister population panels (from *a* and *b*) have 100 haploid genomes each.

**Figure S4.**
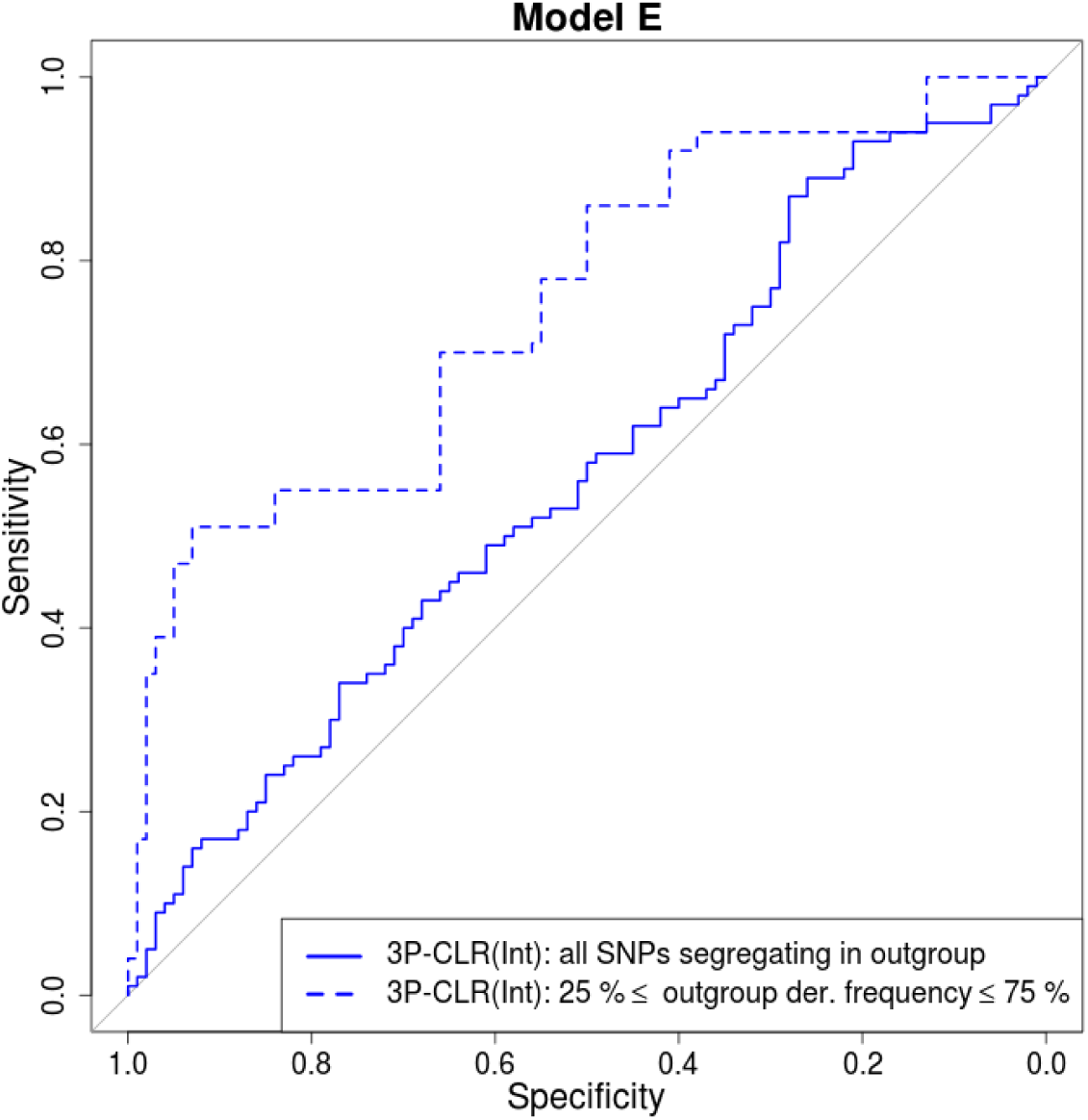
For demographic scenarios with very ancient split times, it is best to use sites segregating at intermediate frequencies in the outgroup. We compared the performance of 3P-CLR(Int) in a demographic scenario with very ancient split times (Model E) under two conditions: including all SNPs that are segregating in the outgroup, and only including SNPs segregating at intermediate frequencies in the outgroup. In both cases, the number of sampled sequences from the outgroup population was 100.

**Figure S5.**
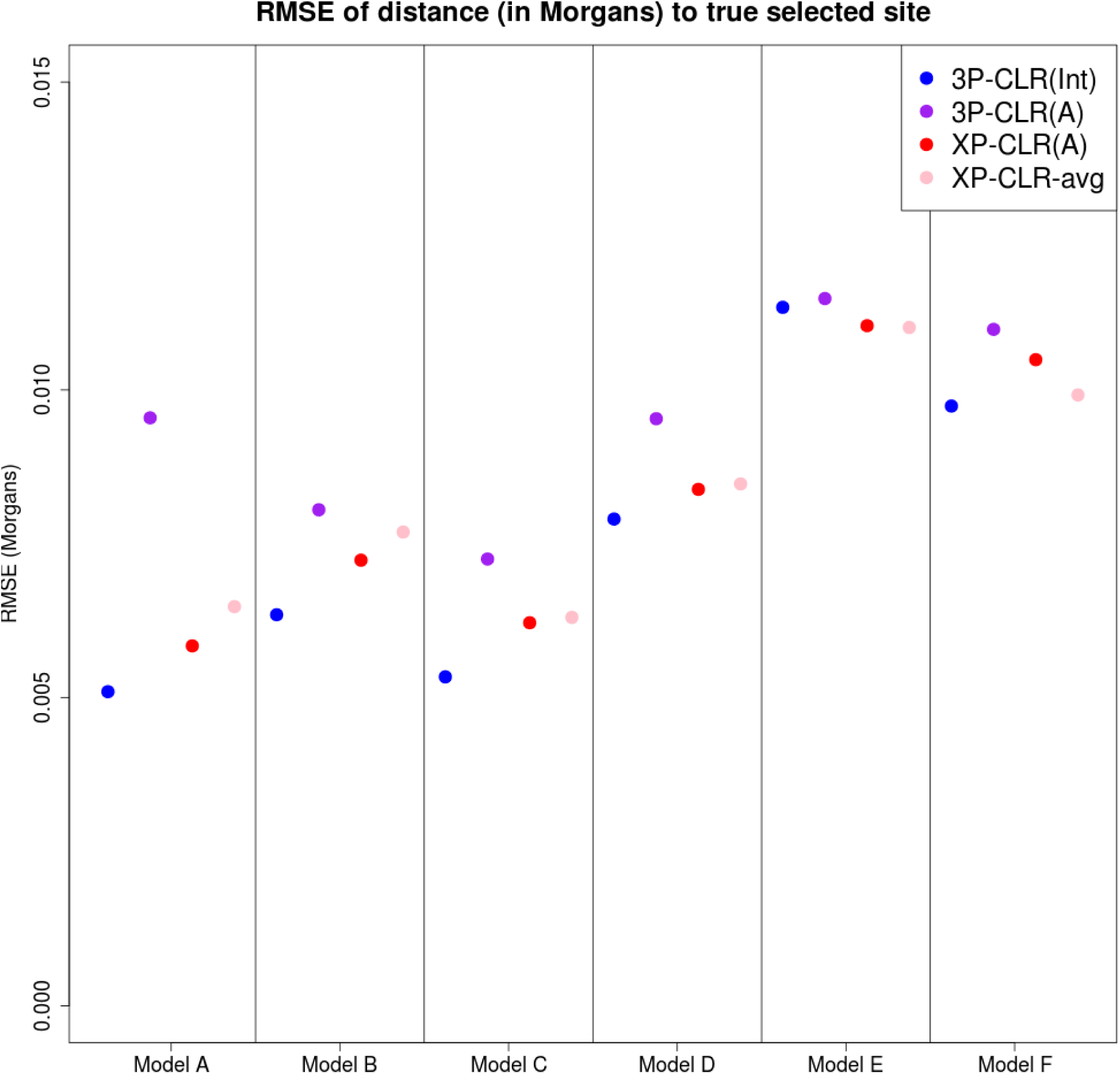
Root-mean squared error for the location of sweeps inferred by 3P-CLR(Int), 3P-CLR(A) and two variants of XP-CLR under different demographic scenarios, when the sweeps occurred before the split of populations *a* and *b*. In this case, the outgroup panel from population *c* contained 100 haploid genomes and the two sister population panels (from *a* and *b*) have 100 haploid genomes each.

**Figure S6.**
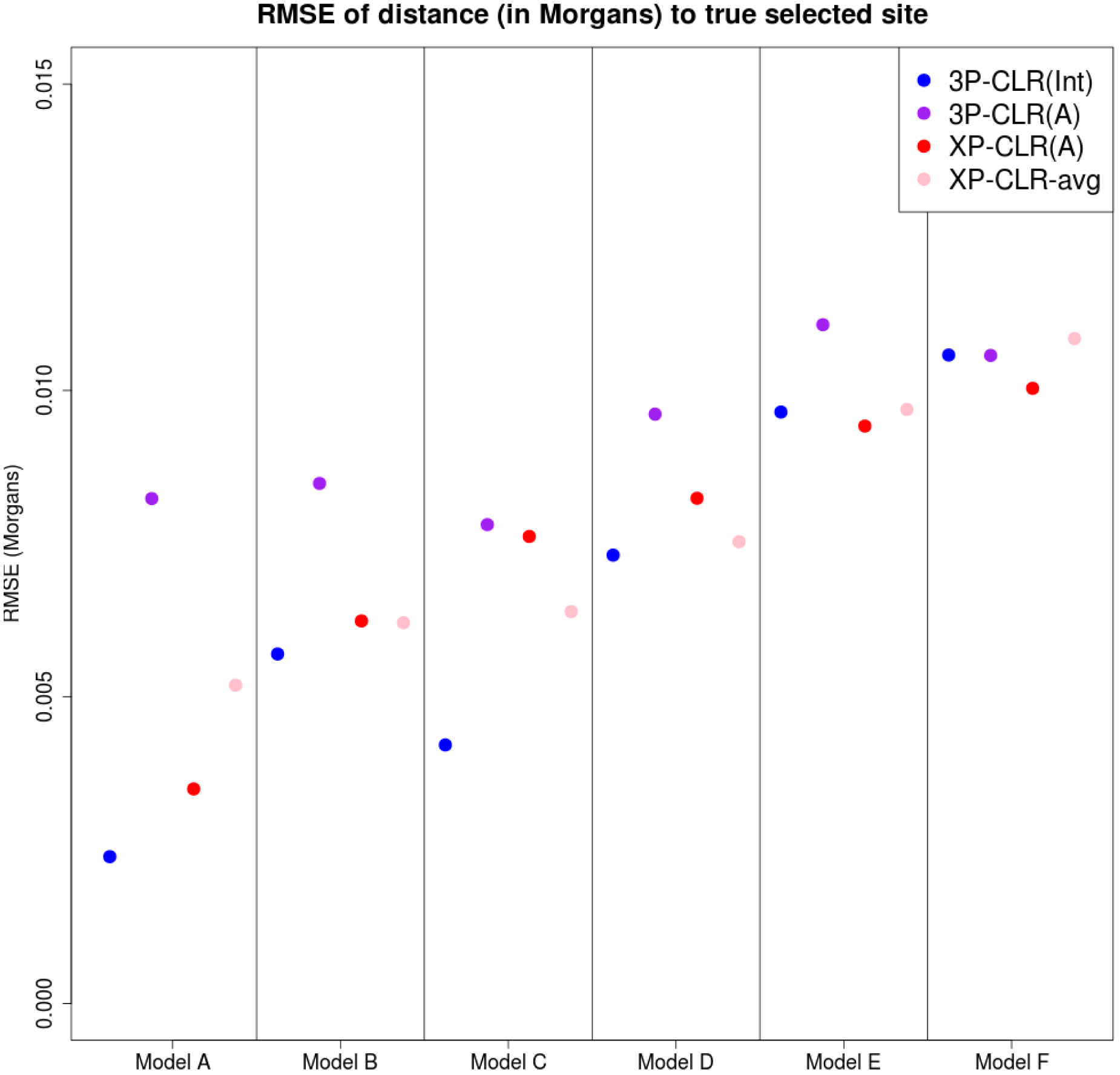
Root-mean squared error for the location of the sweep inferred by 3P-CLR(Int), 3P-CLR(A) and two variants of XP-CLR under different demographic scenarios, when the sweeps occurred before the split of populations *a* and *b*. the outgroup panel from population *c* contained 10 haploid genomes and the two sister population panels (from *a* and *b*) have 100 haploid genomes each.

**Figure S7.**
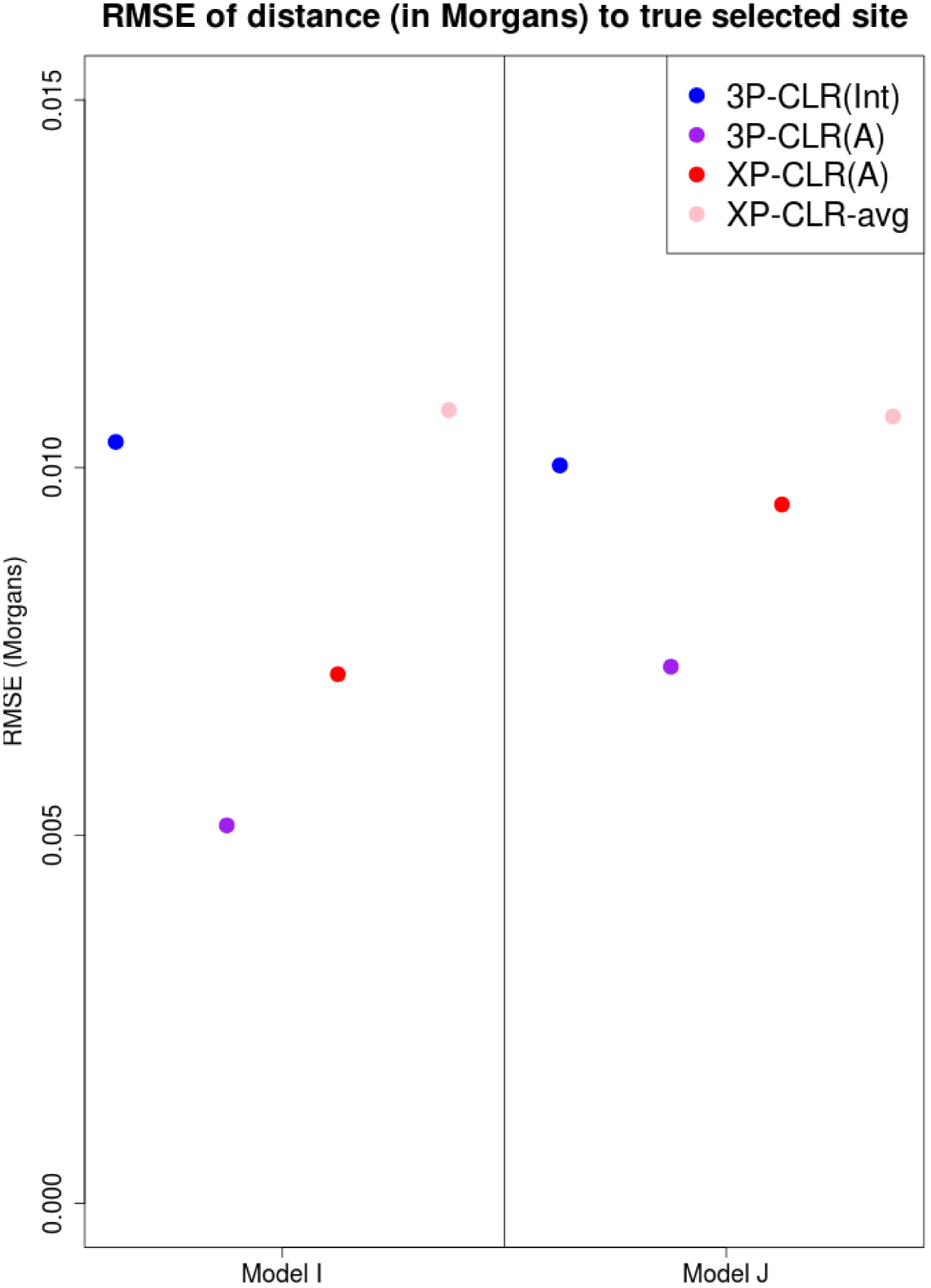
Root-mean squared error for the location of the sweep inferred by 3P-CLR(Int), 3P-CLR(A) and two variants of XP-CLR under different demographic scenarios, when the sweeps occurred in the terminal population branch leading to population *a*, after the split of populations *a* and *b*. In this case, the outgroup panel from population *c* contained 100 haploid genomes and the two sister population panels (from *a* and *b*) have 100 haploid genomes each.

**Figure S8.**
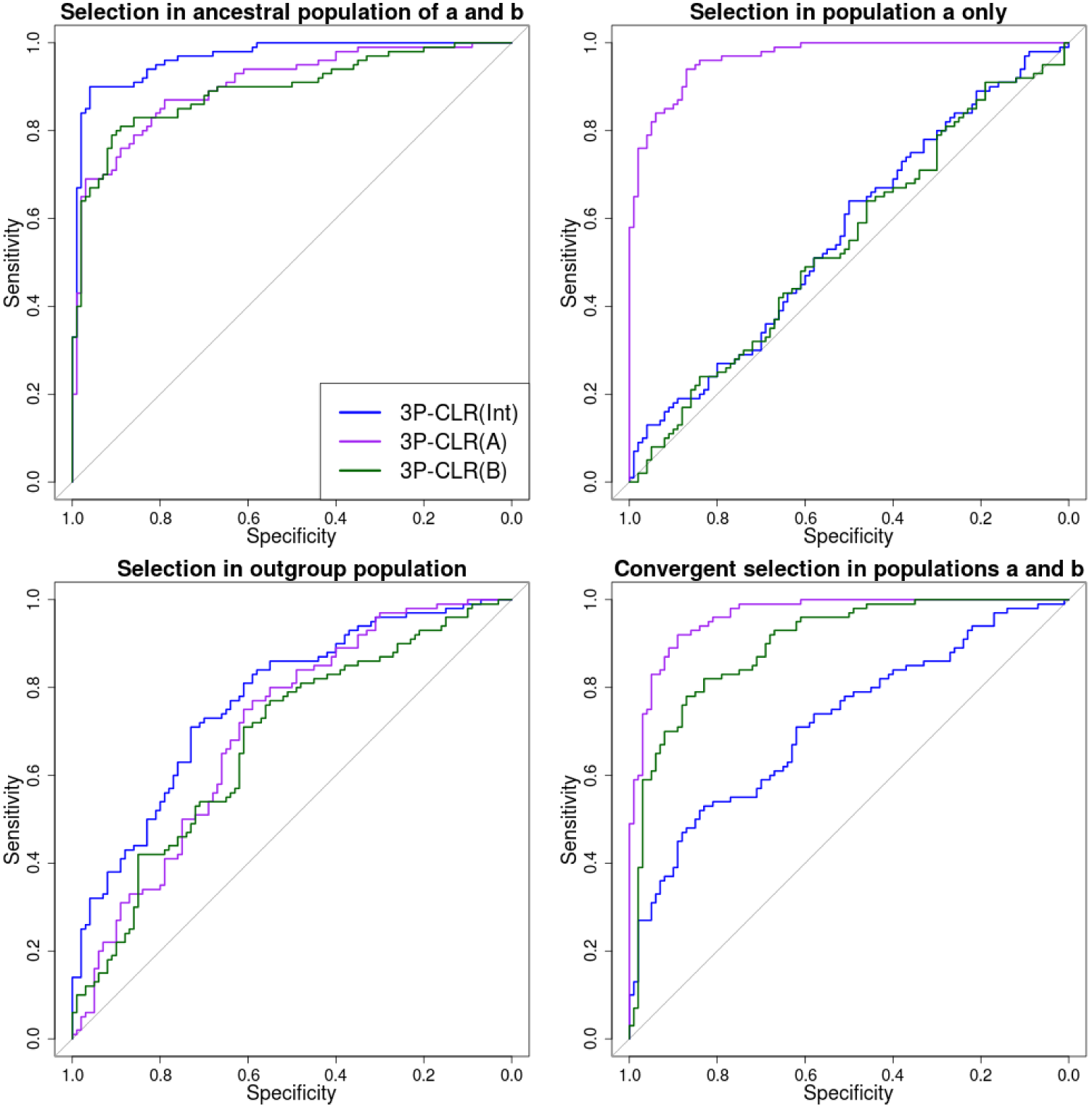
ROC curves for performance of 3P-CLR(Int), 3P-CLR(A) and 3P-CLR(B) when the selective events occur in different branches of the 3-population tree. Upper-left panel: Selection in the ancestral population of populations *a* and *b*. This is the type of events that 3P-CLR(Int) is designed to detect and, therefore, 3P-CLR(Int) is the most sensitive test in this case, though 3P-CLR(A) and 3P-CLR(B) show some sensitivity to these events too. Upper-right panel: Selection exclusive to population *a*. This is the type of events that 3P-CLR(A) is designed to detect, and it is therefore the best-performing statistic in that case, while 3P-CLR(B) and 3P-CLR(Int) are insensitive to selection. Lower-left panel: Selection in the outgroup population. In this case, none of the statistics seem very sensitive to the event, though 3P-CLR(Int) shows better relative sensitivity than the other two statistics. Lower-right panel: Independent selective events in populations *a* and *b* at the same locus. Here, both 3P-CLR(A) and 3P-CLR(B) perform best. In all cases, we used the split times and population sizes specified for Model C.

**Figure S9.**
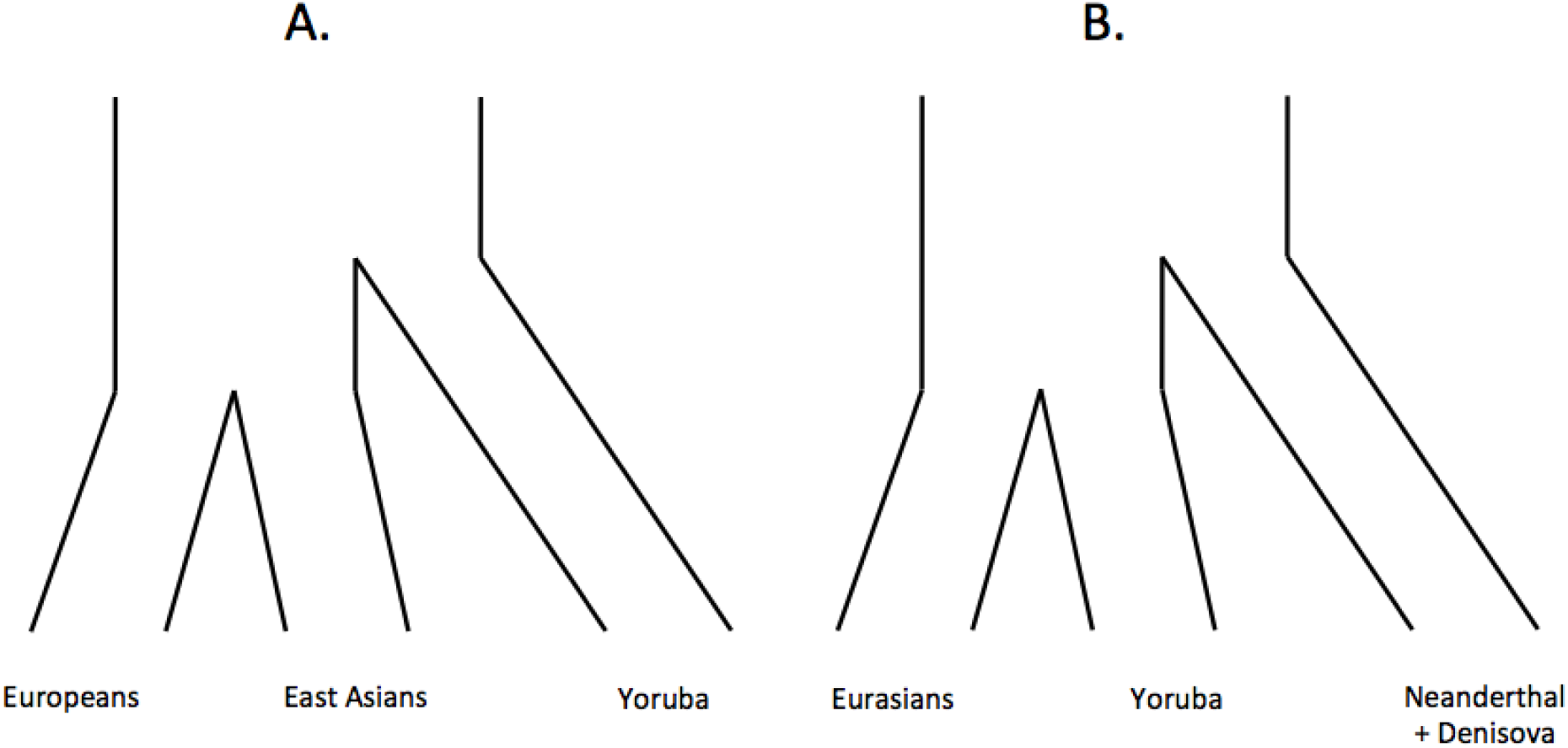
A. Three-population tree separating Europeans, East Asians and Yoruba. B. Three-population tree separating Eurasians, Yoruba and archaic humans (Neanderthal+Denisova).

**Figure S10.**
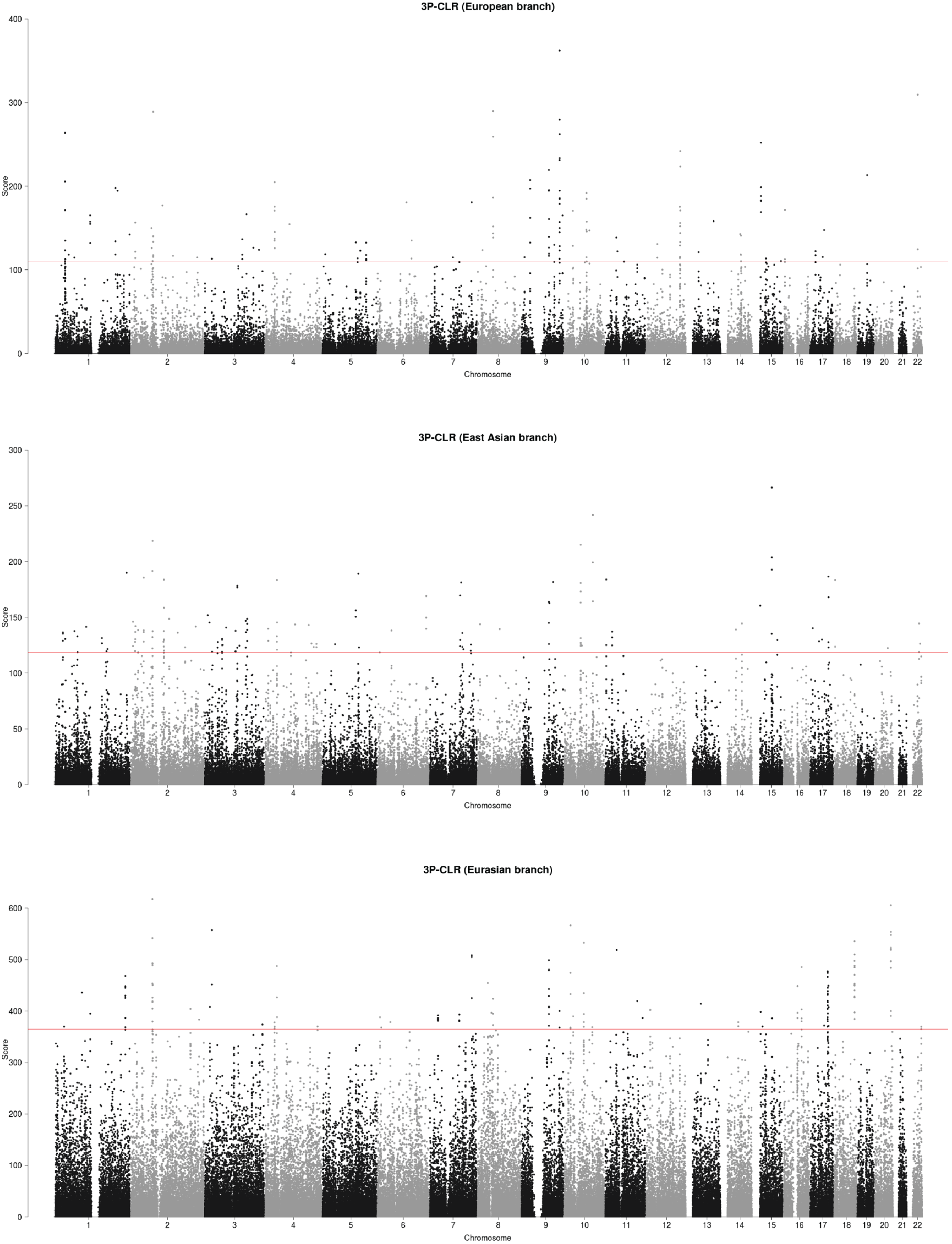
3P-CLR scan of Europeans (upper panel), East Asians (middle panel) and the ancestral population to Europeans and East Asians (lower panel), using Yoruba as the outgroup in all 3 cases. The red line denotes the 99.9% quantile cutoff.

**Figure S11.**
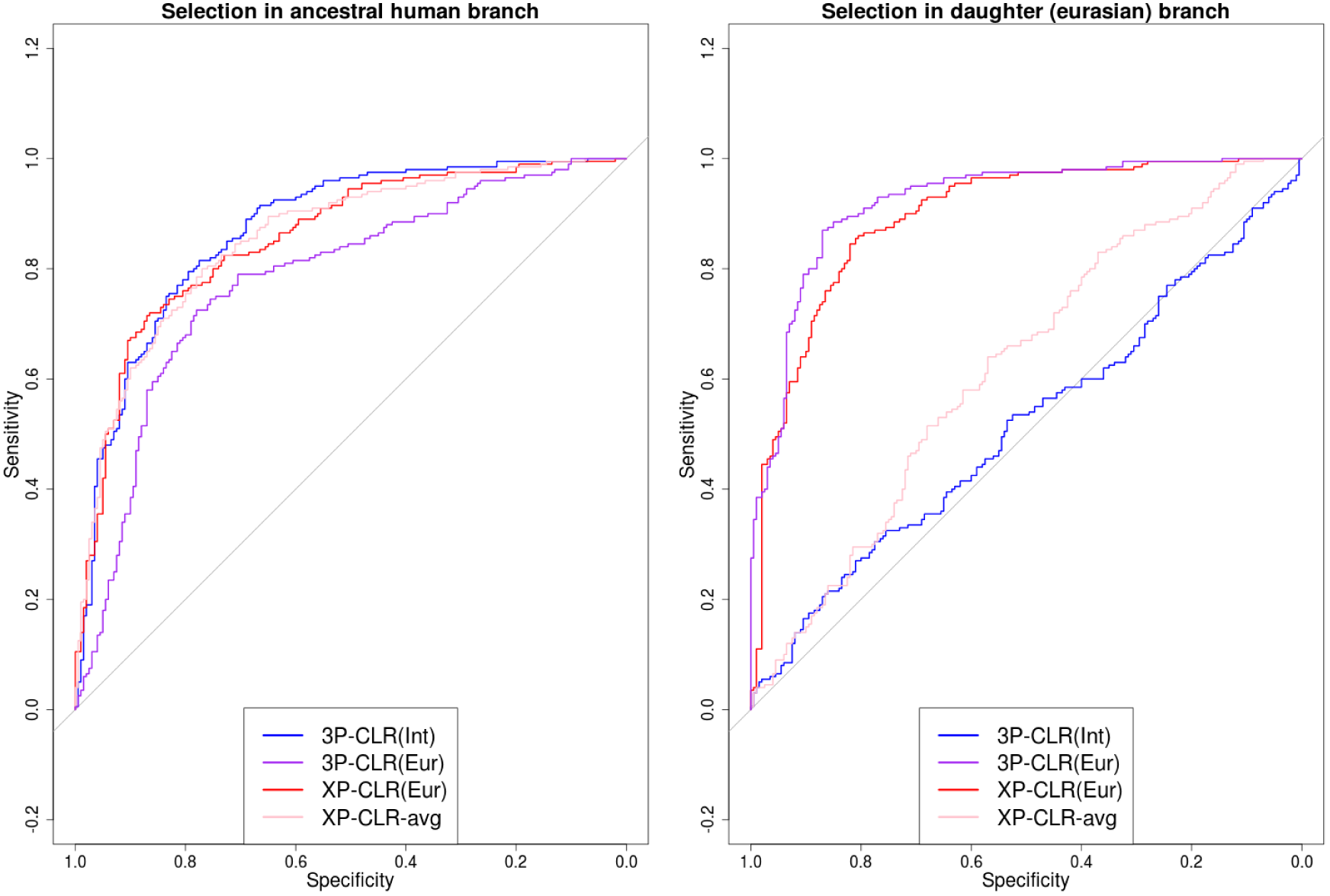
ROC curves for 3P-CLR run to detect selective events in the modern human ancestral branch, using simulations incorporating the history of population size changes and Neanderthal-to-Eurasian admixture inferred in Prüfer et al. (2014).

**Figure S12.**
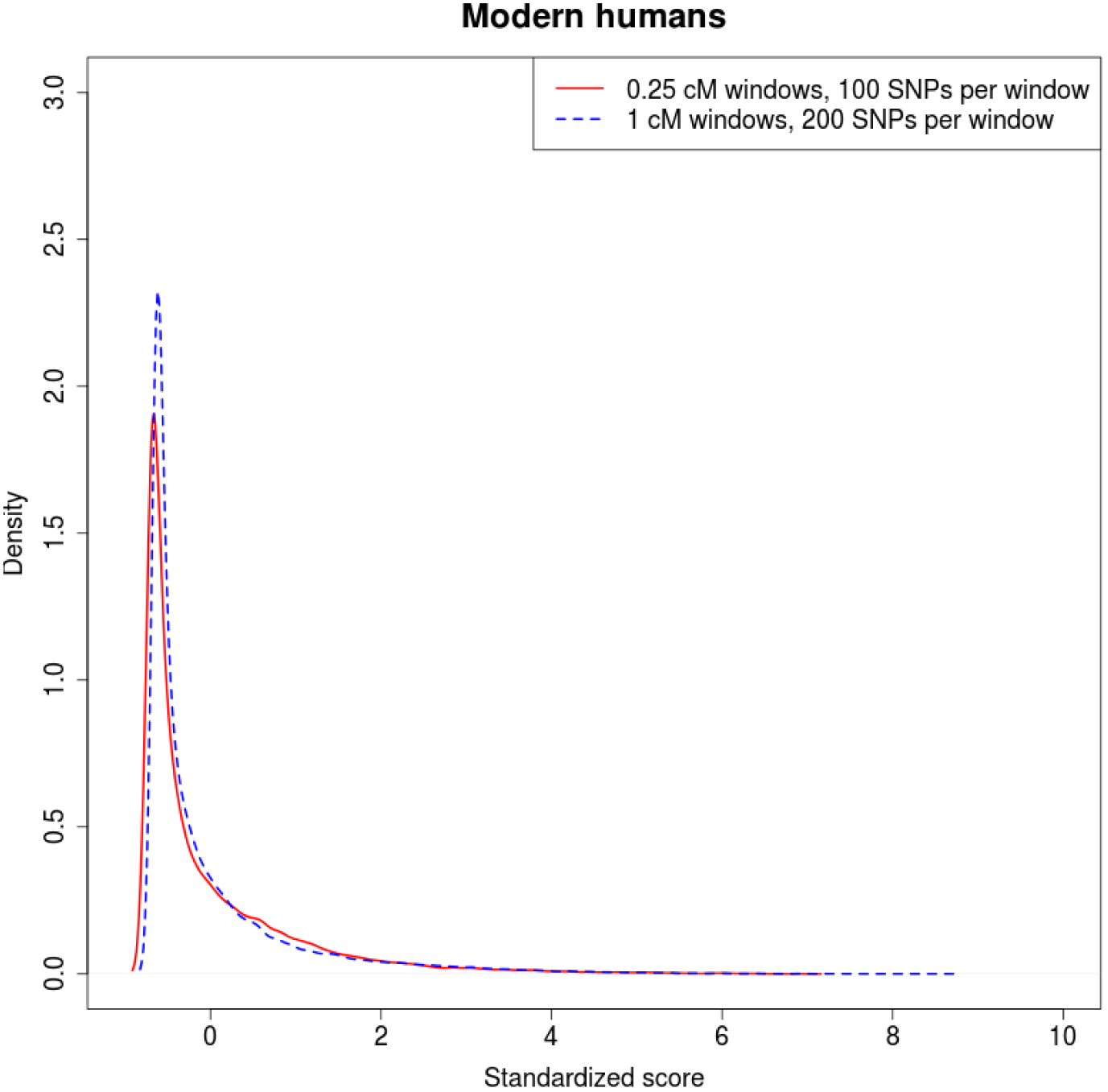
Comparison of 3P-CLR on the modern human ancestral branch under different window sizes and central SNP spacing. The red density is the density of standardized scores for 3P-CLR run using 0.25 cM windows, 100 SNPs per window and a spacing of 10 SNPs between each central SNP. The blue dashed density is the density of standardized scores for 3P-CLR run using 1 cM windows, 200 SNPs per window and a spacing of 40 SNPs between each central SNP.

**Figure S13.**
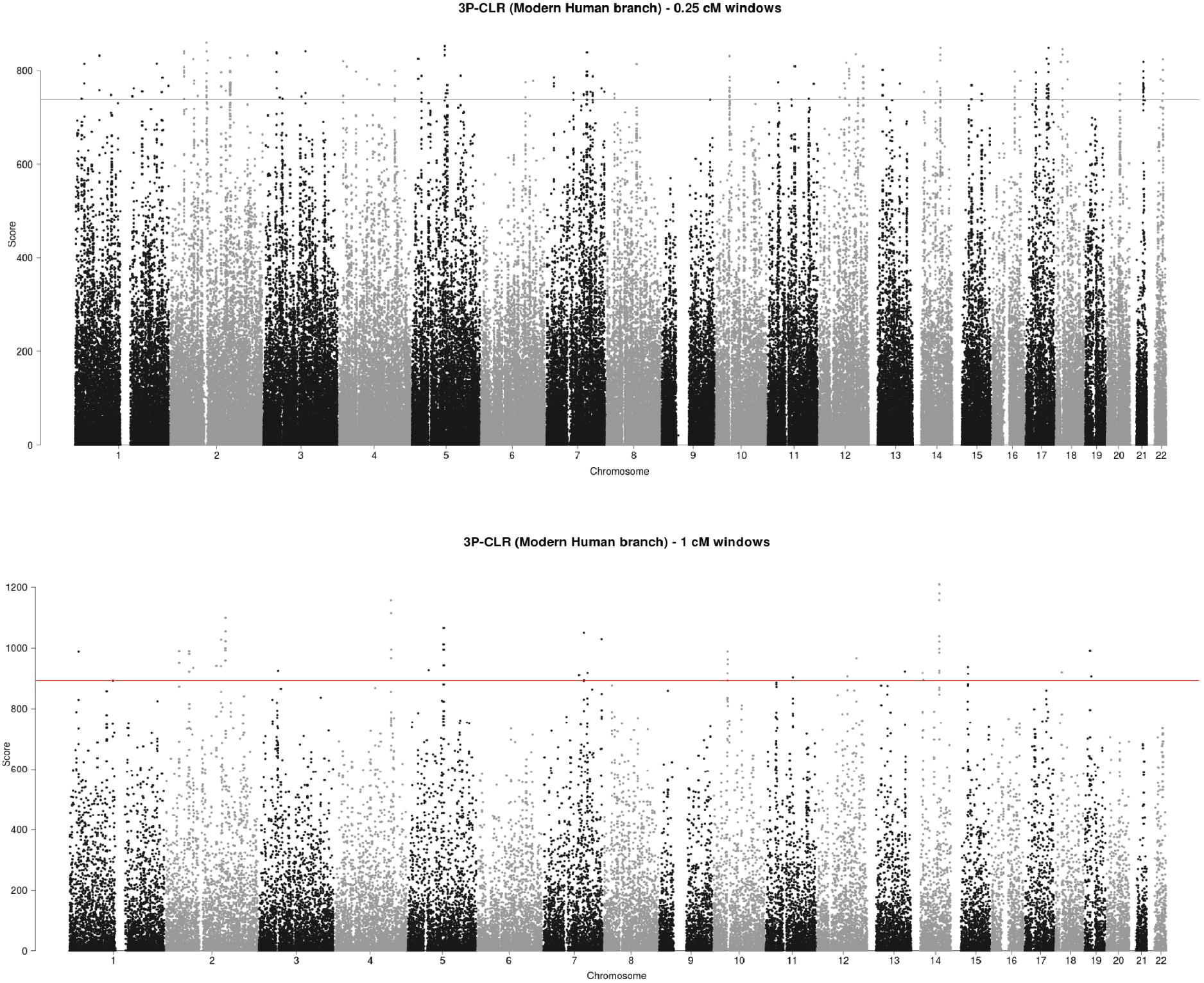
3P-CLR scan of the ancestral branch to Yoruba and Eurasians, using the Denisovan and Neanderthal genomes as the outgroup. The red line denotes the 99.9% quantile cutoff. The top panel shows a run using 0.25 cM windows, each containing 100 SNPs, and sampling a candidate beneficial SNP every 10 SNPs. The bottom panels shows a run using 1 cM windows, each containing 200 SNPs, and sampling a candidate beneficial SNP every 40 SNPs.

**Figure S14.**
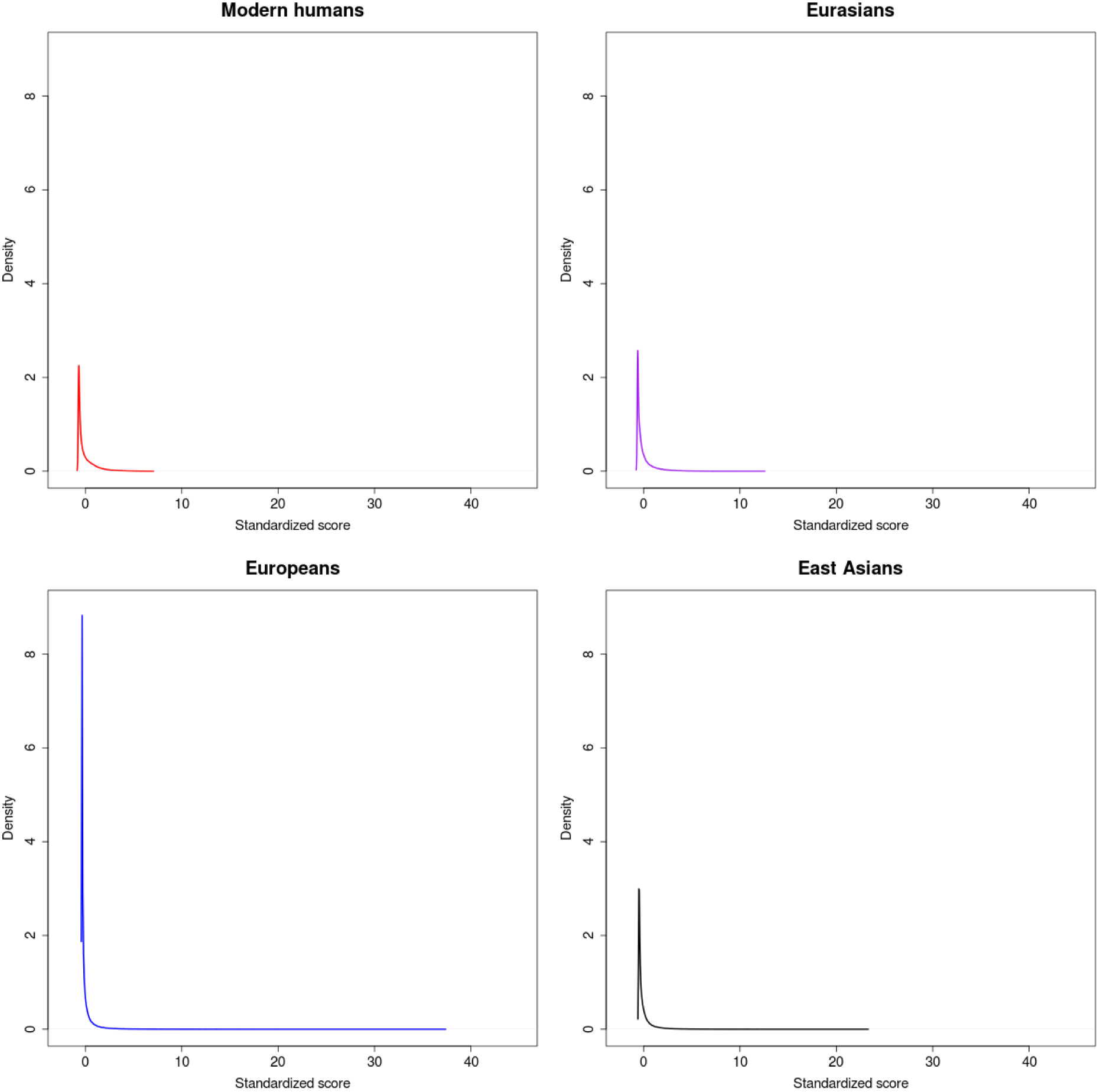
Genome-wide densities of each of the 3P-CLR scores described in this work. The distributions of scores testing for recent selection (Europeans and East Asians) have much longer tails than the distributions of scores testing for more ancient selection (Modern Humans and Eurasians). All scores were computed using 0.25 cM windows and were then standardized using their genome-wide means and standard deviations.

**Figure S15.**
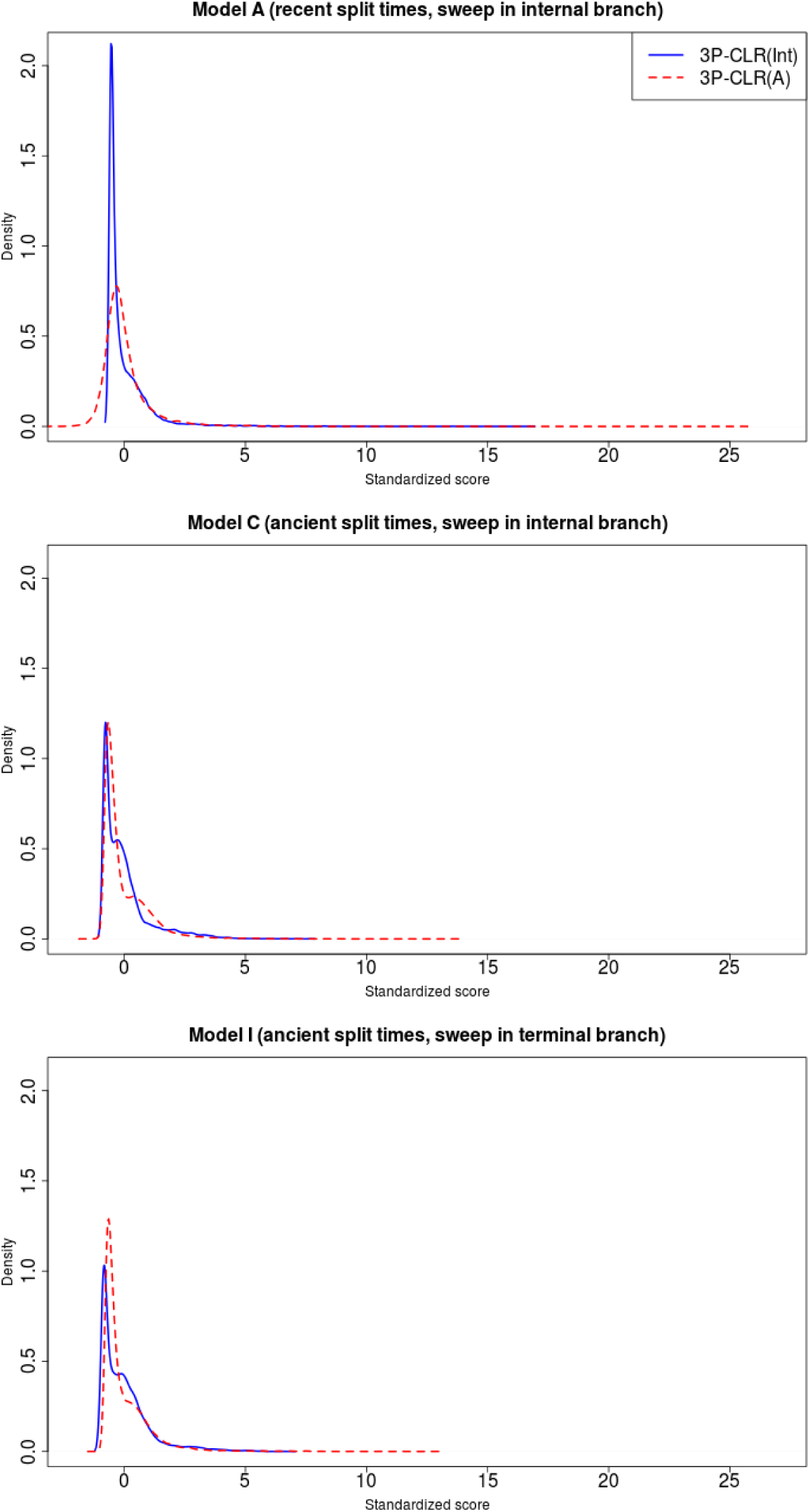
Distribution of 3P-CLR(Int) and 3P-CLR(A) scores under different demographic histories. We combined all scores obtained from 100 neutral simulations and 100 simulations with a selective sweep under different demographic and selection regimes. We then plotted the densities of the resulting scores. Top panel: Model A; Middle panel: Model C; Bottom panel: Model I. See Table 1 for details about each model.

## Literature Cited

Goncalo R Abecasis, Adam Auton, Lisa D Brooks, Mark A DePristo, Richard M Durbin, Robert E Handsaker, Hyun Min Kang, Gabor T Marth, Gil A McVean, Hans Lehrach, et al. An integrated map of genetic variation from 1,092 human genomes. Nature, 491(7422):56–65, 2012.

Joshua M Akey, Ge Zhang, Kun Zhang, Li Jin, and Mark D Shriver. Interrogating a high-density snp map for signatures of natural selection. Genome research, 12(12):1805–1814, 2002.

Francesca Ariani, Giuseppe Hayek, Dalila Rondinella, Rosangela Artuso, Maria Antonietta Mencarelli, Ariele Spanhol-Rosseto, Marzia Pollazzon, Sabrina Buoni, Ottavia Spiga, Sara Ricciardi, et al. Foxg1 is responsible for the congenital variant of rett syndrome. The American Journal of Human Genetics, 83(1):89–93, 2008.

Wojciech Branicki, Urszula Brudnik, and Anna Wojas-Pelc. Interactions between herc2, oca2 and mc1r may influence human pigmentation phenotype. Annals of human genetics, 73(2):160–170, 2009.

David Brawand, Magali Soumillon, Anamaria Necsulea, Philippe Julien, Gábor Csárdi, Patrick Harrigan, Manuela Weier, Angélica Liechti, Ayinuer Aximu-Petri, Martin Kircher, et al. The evolution of gene expression levels in mammalian organs. Nature, 478(7369):343–348, 2011.

Sergi Castellano, Genís Parra, Federico A Sánchez-Quinto, Fernando Racimo, Martin Kuhlwilm, Martin Kircher, Susanna Sawyer, Qiaomei Fu, Anja Heinze, Birgit Nickel, et al. Patterns of coding variation in the complete exomes of three neandertals. Proceedings of the National Academy of Sciences, 111(18):6666–6671, 2014.

Hua Chen, Nick Patterson, and David Reich. Population differentiation as a test for selective sweeps. Genome research, 20(3):393–402, 2010.

Gregory M Cooper, David L Goode, Sarah B Ng, Arend Sidow, Michael J Bamshad, Jay Shendure, and Deborah A Nickerson. Single-nucleotide evolutionary constraint scores highlight disease-causing mutations. Nature methods, 7(4):250–251, 2010.

Jessica L Crisci, Alex Wong, Jeffrey M Good, and Jeffrey D Jensen. On characterizing adaptive events unique to modern humans. Genome biology and evolution, 3:791–798, 2011.

Kevin L Du, Mary Chen, Jian Li, John J Lepore, Patricia Mericko, and Michael S Parmacek. Megakaryoblastic leukemia factor-1 transduces cytoskeletal signals and induces smooth muscle cell differentiation from undifferentiated embryonic stem cells. Journal of Biological Chemistry, 279(17):17578–17586, 2004.

I Dunham, A Kundaje, SF Aldred, PJ Collins, C Davis, F Doyle, CB Epstein, S Frietze, J Harrow, R Kaul, et al. An integrated encyclopedia of dna elements in the human genome. Nature, 489(7414): 57–74, 2012.

Richard Durrett and Jason Schweinsberg. Approximating selective sweeps. Theoretical population biology, 66(2):129–138, 2004.

Howard J Edenberg, Danielle M Dick, Xiaoling Xuei, Huijun Tian, Laura Almasy, Lance O Bauer, Raymond R Crowe, Alison Goate, Victor Hesselbrock, Kevin Jones, et al. Variations in gabra2, encoding the *α*2 subunit of the gaba a receptor, are associated with alcohol dependence and with brain oscillations. The American Journal of Human Genetics, 74(4):705–714, 2004.

Patrick Edery, Stéphane Chabrier, Irène Ceballos-Picot, Sandrine Marie, Marie-Françoise Vincent, and Marc Tardieu. Intrafamilial variability in the phenotypic expression of adenylosuccinate lyase deficiency: a report on three patients. American Journal of Medical Genetics Part A, 120(2):185–190, 2003.

Iiro Eerola, Laurence M Boon, John B Mulliken, Patricia E Burrows, Anne Dompmartin, Shoji Watanabe, Romain Vanwijck, and Miikka Vikkula. Capillary malformation–arteriovenous malformation, a new clinical and genetic disorder caused by rasa1 mutations. The American Journal of Human Genetics, 73(6):1240–1249, 2003.

Hans Eiberg, Jesper Troelsen, Mette Nielsen, Annemette Mikkelsen, Jonas Mengel-From, Klaus W Kjaer, and Lars Hansen. Blue eye color in humans may be caused by a perfectly associated founder mutation in a regulatory element located within the herc2 gene inhibiting oca2 expression. Human genetics, 123(2):177–187, 2008.

Warren J Ewens. Mathematical Population Genetics 1: Theoretical Introduction, volume 27. Springer Science & Business Media, 2012.

María Inés Fariello, Simon Boitard, Hugo Naya, Magali SanCristobal, and Bertrand Servin. Detecting signatures of selection through haplotype differentiation among hierarchically structured populations. Genetics, 193(3):929–941, 2013.

Justin C Fay and Chung-I Wu. Hitchhiking under positive darwinian selection. Genetics, 155(3):1405–1413, 2000.

Joseph Felsenstein. Evolutionary trees from gene frequencies and quantitative characters: finding maximum likelihood estimates. Evolution, pages 1229–1242, 1981.

Eitan Friedman, Pablo V Gejman, George A Martin, and Frank McCormick. Nonsense mutations in the c–terminal sh2 region of the gtpase activating protein (gap) gene in human tumours. Nature genetics, 5(3):242–247, 1993.

Qiaomei Fu, Heng Li, Priya Moorjani, Flora Jay, Sergey M Slepchenko, Aleksei A Bondarev, Philip LF Johnson, Ayinuer Aximu-Petri, Kay Prüfer, Cesare de Filippo, et al. Genome sequence of a 45,000-year-old modern human from western siberia. Nature, 514(7523):445–449, 2014.

Akihiro Fujimoto, Ryosuke Kimura, Jun Ohashi, Kazuya Omi, Rika Yuliwulandari, Lilian Batubara, Mohammad Syamsul Mustofa, Urai Samakkarn, Wannapa Settheetham-Ishida, Takafumi Ishida, et al. A scan for genetic determinants of human hair morphology: Edar is associated with asian hair thickness. Human Molecular Genetics, 17(6):835–843, 2008.

Qingshen Gao, Seetha Srinivasan, Sarah N Boyer, David E Wazer, and Vimla Band. The e6 oncoproteins of high-risk papillomaviruses bind to a novel putative gap protein, e6tp1, and target it for degradation. Molecular and cellular biology, 19(1):733–744, 1999.

Cyril Gitiaux, Irène Ceballos-Picot, Sandrine Marie, Vassili Valayannopoulos, Marlène Rio, Séverine Verrieres, Jean François Benoist, Marie Françoise Vincent, Isabelle Desguerre, and Nadia Bahi-Buisson. Misleading behavioural phenotype with adenylosuccinate lyase deficiency. European Journal of Human Genetics, 17(1):133–136, 2009.

Shiaoching Gong, Chen Zheng, Martin L Doughty, Kasia Losos, Nicholas Didkovsky, Uta B Schambra, Norma J Nowak, Alexandra Joyner, Gabrielle Leblanc, Mary E Hatten, et al. A gene expression atlas of the central nervous system based on bacterial artificial chromosomes. Nature, 425(6961):917–925, 2003.

Richard E Green, Johannes Krause, Adrian W Briggs, Tomislav Maricic, Udo Stenzel, Martin Kircher, Nick Patterson, Heng Li, Weiwei Zhai, Markus Hsi-Yang Fritz, et al. A draft sequence of the neandertal genome. science, 328(5979):710–722, 2010.

Sharon R Grossman, Ilya Shylakhter, Elinor K Karlsson, Elizabeth H Byrne, Shannon Morales, Gabriel Frieden, Elizabeth Hostetter, Elaine Angelino, Manuel Garber, Or Zuk, et al. A composite of multiple signals distinguishes causal variants in regions of positive selection. Science, 327(5967):883–886, 2010.

Julius Gudmundsson, Patrick Sulem, Thorunn Rafnar, Jon T Bergthorsson, Andrei Manolescu, Daniel Gudbjartsson, Bjarni A Agnarsson, Asgeir Sigurdsson, Kristrun R Benediktsdottir, Thorarinn Blondal, et al. Common sequence variants on 2p15 and xp11. 22 confer susceptibility to prostate cancer. Nature genetics, 40(3):281–283, 2008.

Adilson Guilherme, Neil A Soriano, Paul S Furcinitti, and Michael P Czech. Role of ehd1 and ehbp1 in perinuclear sorting and insulin-regulated glut4 recycling in 3t3-l1 adipocytes. Journal of Biological Chemistry, 279(38):40062–40075, 2004.

Ryan N Gutenkunst, Ryan D Hernandez, Scott H Williamson, and Carlos D Bustamante. Inferring the joint demographic history of multiple populations from multidimensional snp frequency data. PLoS genetics, 5(10):e1000695, 2009.

Ruth Halaban and Gisela Moellmann. Murine and human b locus pigmentation genes encode a glycoprotein (gp75) with catalase activity. Proceedings of the National Academy of Sciences, 87(12):4809–4813, 1990.

Jiali Han, Peter Kraft, Hongmei Nan, Qun Guo, Constance Chen, Abrar Qureshi, Susan E Hankinson, Frank B Hu, David L Duffy, Zhen Zhen Zhao, et al. A genome-wide association study identifies novel alleles associated with hair color and skin pigmentation. PLoS genetics, 4(5):e1000074, 2008.

Marc Henrion, Matthew Frampton, Ghislaine Scelo, Mark Purdue, Yuanqing Ye, Peter Broderick, Alastair Ritchie, Richard Kaplan, Angela Meade, James McKay, et al. Common variation at 2q22. 3 (zeb2) influences the risk of renal cancer. Human molecular genetics, 22(4):825–831, 2013.

Ryan D Hernandez, Joanna L Kelley, Eyal Elyashiv, S Cord Melton, Adam Auton, Gilean McVean, Guy Sella, Molly Przeworski, et al. Classic selective sweeps were rare in recent human evolution. science, 331(6019):920–924, 2011.

D Hershkovitz, D Bercovich, E Sprecher, and M Lapidot. Rasa1 mutations may cause hereditary capillary malformations without arteriovenous malformations. British Journal of Dermatology, 158(5):1035–1040, 2008.

Anjali G Hinch, Arti Tandon, Nick Patterson, Yunli Song, Nadin Rohland, Cameron D Palmer, Gary K Chen, Kai Wang, Sarah G Buxbaum, Ermeg L Akylbekova, et al. The landscape of recombination in african americans. Nature, 476(7359):170–175, 2011.

Karen A Hunt, Alexandra Zhernakova, Graham Turner, Graham AR Heap, Lude Franke, Marcel Bruinenberg, Jihane Romanos, Lotte C Dinesen, Anthony W Ryan, Davinder Panesar, et al. Newly identified genetic risk variants for celiac disease related to the immune response. Nature genetics, 40(4):395–402, 2008.

Minoru Kanehisa and Susumu Goto. Kegg: kyoto encyclopedia of genes and genomes. Nucleic acids research, 28(1):27–30, 2000.

Eimear E Kenny, Nicholas J Timpson, Martin Sikora, Muh-Ching Yee, Andrés Moreno-Estrada, Celeste Eng, Scott Huntsman, Esteban González Burchard, Mark Stoneking, Carlos D Bustamante, et al. Melanesian blond hair is caused by an amino acid change in tyrp1. Science, 336(6081):554–554, 2012.

Ryosuke Kimura, Tetsutaro Yamaguchi, Mayako Takeda, Osamu Kondo, Takashi Toma, Kuniaki Haneji, Tsunehiko Hanihara, Hirotaka Matsukusa, Shoji Kawamura, Koutaro Maki, et al. A common variation in edar is a genetic determinant of shovel-shaped incisors. The American Journal of Human Genetics, 85(4):528–535, 2009.

Martin Kircher, Daniela M Witten, Preti Jain, Brian J O’Roak, Gregory M Cooper, and Jay Shendure. A general framework for estimating the relative pathogenicity of human genetic variants. Nature genetics, 46(3):310–315, 2014.

Stanislav Kmoch, Hana Hartmannová, Blanka Stibùrková, Jakub Krijt, Marie Zikánová, and Ivan Šebesta. Human adenylosuccinate lyase (adsl), cloning and characterization of full-length cdna and its isoform, gene structure and molecular basis for adsl deficiency in six patients. Human molecular genetics, 9(10):1501–1513, 2000.

Julia Knabl, Robert Witschi, Katharina Hösl, Heiko Reinold, Ulrike B Zeilhofer, Seifollah Ahmadi, Johannes Brockhaus, Marina Sergejeva, Andreas Hess, Kay Brune, et al. Reversal of pathological pain through specific spinal gabaa receptor subtypes. Nature, 451(7176):330–334, 2008.

Robert Kofler and Christian Schlötterer. Gowinda: unbiased analysis of gene set enrichment for genome-wide association studies. Bioinformatics, 28(15):2084–2085, 2012.

Iosif Lazaridis, Nick Patterson, Alissa Mittnik, Gabriel Renaud, Swapan Mallick, Karola Kirsanow, Peter H Sudmant, Joshua G Schraiber, Sergi Castellano, Mark Lipson, et al. Ancient human genomes suggest three ancestral populations for present-day europeans. Nature, 513(7518):409–413, 2014.

RC Lewontin and Jesse Krakauer. Distribution of gene frequency as a test of the theory of the selective neutrality of polymorphisms. Genetics, 74(1):175–195, 1973.

Heng Li and Richard Durbin. Inference of human population history from individual whole-genome sequences. Nature, 475(7357):493–496, 2011.

Mulin Jun Li, Panwen Wang, Xiaorong Liu, Ee Lyn Lim, Zhangyong Wang, Meredith Yeager, Maria P Wong, Pak Chung Sham, Stephen J Chanock, and Junwen Wang. Gwasdb: a database for human genetic variants identified by genome-wide association studies. Nucleic acids research, page gkr1182, 2011.

Bruce G Lindsay. Composite likelihood methods. Contemporary Mathematics, 80(1):221–39, 1988.

Mark Lipson, Po-Ru Loh, Alex Levin, David Reich, Nick Patterson, and Bonnie Berger. Efficient moment-based inference of admixture parameters and sources of gene flow. Molecular biology and evolution, 30(8):1788–1802, 2013.

PD Maaswinkel-Mooij, LAEM Laan, W Onkenhout, OF Brouwer, J Jaeken, and BJHM Poorthuis. Adenylosuccinase deficiency presenting with epilepsy in early infancy. Journal of inherited metabolic disease, 20(4):606–607, 1997.

Sandrine Marie, Harry Cuppens, Michel Heuterspreute, Martine Jaspers, Eduardo Zambrano Tola, Xiao Xiao Gu, Eric Legius, M Vincent, Jaak Jaeken, Jean-Jacques Cassiman, et al. Mutation analysis in adenylosuccinate lyase deficiency: Eight novel mutations in the re-evaluated full adsl coding sequence. Human mutation, 13(3):197–202, 1999.

Iain Mathieson, Iosif Lazaridis, Nadin Rohland, Swapan Mallick, Bastien Llamas, Joseph Pickrell, Harald Meller, Manuel A Rojo Guerra, Johannes Krause, David Anthony, et al. Eight thousand years of natural selection in europe. bioRxiv, page 016477, 2015.

Gunter Meister, Markus Landthaler, Lasse Peters, Po Yu Chen, Henning Urlaub, Reinhard Lührmann, and Thomas Tuschl. Identification of novel argonaute-associated proteins. Current biology, 15(23):2149–2155, 2005.

MA Mencarelli, A Spanhol-Rosseto, R Artuso, D Rondinella, R De Filippis, N Bahi-Buisson, J Nectoux, R Rubinsztajn, T Bienvenu, A Moncla, et al. Novel foxg1 mutations associated with the congenital variant of rett syndrome. Journal of medical genetics, 47(1):49–53, 2010.

Thomas Mercher, Maryvonne Busson-Le Coniat, Richard Monni, Martine Mauchauffé, Florence Nguyen Khac, Lætitia Gressin, Francine Mugneret, Thierry Leblanc, Nicole Dastugue, Roland Berger, et al. Involvement of a human gene related to the drosophila spen gene in the recurrent t (1; 22) translocation of acute megakaryocytic leukemia. Proceedings of the National Academy of Sciences, 98(10):5776–5779, 2001.

Philipp W Messer. Slim: simulating evolution with selection and linkage. Genetics, 194(4):1037–1039, 2013.

Matthias Meyer, Martin Kircher, Marie-Theres Gansauge, Heng Li, Fernando Racimo, Swapan Mallick, Joshua G Schraiber, Flora Jay, Kay Prüfer, Cesare de Filippo, et al. A high-coverage genome sequence from an archaic denisovan individual. Science, 338(6104):222–226, 2012.

George Nicholson, Albert V Smith, Frosti Jónsson, Ómar Gústafsson, Kári Stefánsson, and Peter Donnelly. Assessing population differentiation and isolation from single-nucleotide polymorphism data. Journal of the Royal Statistical Society: Series B (Statistical Methodology), 64(4):695–715, 2002.

Taras K Oleksyk, Kai Zhao, M Francisco, Dennis A Gilbert, Stephen J O’Brien, and Michael W Smith. Identifying selected regions from heterozygosity and divergence using a light-coverage genomic dataset from two human populations. PLoS One, 3(3):e1712, 2008.

Luigi Pace, Alessandra Salvan, and Nicola Sartori. Adjusting composite likelihood ratio statistics. Statistica Sinica, 21(1):129, 2011.

Lavinia Paternoster, David M Evans, Ellen Aagaard Nohr, Claus Holst, Valerie Gaborieau, Paul Brennan, Anette Prior Gjesing, Niels Grarup, Daniel R Witte, Torben Jørgensen, et al. Genome-wide population-based association study of extremely overweight young adults–the goya study. PLoS One, 6(9):e24303, 2011.

Nick Patterson, Priya Moorjani, Yontao Luo, Swapan Mallick, Nadin Rohland, Yiping Zhan, Teri Genschoreck, Teresa Webster, and David Reich. Ancient admixture in human history. Genetics, 192(3): 1065–1093, 2012.

Len A Pennacchio, Nadav Ahituv, Alan M Moses, Shyam Prabhakar, Marcelo A Nobrega, Malak Shoukry, Simon Minovitsky, Inna Dubchak, Amy Holt, Keith D Lewis, et al. In vivo enhancer analysis of human conserved non-coding sequences. Nature, 444(7118):499–502, 2006.

Roy H Perlis, Jie Huang, Shaun Purcell, Maurizio Fava, A John Rush, Patrick F Sullivan, Steven P Hamilton, Francis J McMahon, Thomas Schulze, James B Potash, et al. Genome-wide association study of suicide attempts in mood disorder patients. Genome, 167(12), 2010.

Joseph K Pickrell and Jonathan K Pritchard. Inference of population splits and mixtures from genome-wide allele frequency data. PLoS genetics, 8(11):e1002967, 2012.

Joseph K Pickrell, Graham Coop, John Novembre, Sridhar Kudaravalli, Jun Z Li, Devin Absher, Balaji S Srinivasan, Gregory S Barsh, Richard M Myers, Marcus W Feldman, et al. Signals of recent positive selection in a worldwide sample of human populations. Genome research, 19(5):826–837, 2009.

Dimitrina D Pravtcheva and Thomas L Wise. Disruption of apc10/doc1 in three alleles of oligosyn-dactylism. Genomics, 72(1):78–87, 2001.

Kay Prüfer, Fernando Racimo, Nick Patterson, Flora Jay, Sriram Sankararaman, Susanna Sawyer, Anja Heinze, Gabriel Renaud, Peter H Sudmant, Cesare de Filippo, et al. The complete genome sequence of a neanderthal from the altai mountains. Nature, 505(7481):43–49, 2014.

Valérie Race, Sandrine Marie, Marie-Françoise Vincent, and Georges Van den Berghe. Clinical, biochemical and molecular genetic correlations in adenylosuccinate lyase deficiency. Human molecular genetics, 9(14):2159–2165, 2000.

Fernando Racimo, Martin Kuhlwilm, and Montgomery Slatkin. A test for ancient selective sweeps and an application to candidate sites in modern humans. Molecular biology and evolution, 31(12):3344–3358, 2014.

Peter N Robinson, Sebastian Köhler, Sebastian Bauer, Dominik Seelow, Denise Horn, and Stefan Mundlos. The human phenotype ontology: a tool for annotating and analyzing human hereditary disease. The American Journal of Human Genetics, 83(5):610–615, 2008.

Kate R Rosenbloom, Timothy R Dreszer, Jeffrey C Long, Venkat S Malladi, Cricket A Sloan, Brian J Raney, Melissa S Cline, Donna Karolchik, Galt P Barber, Hiram Clawson, et al. Encode whole-genome data in the ucsc genome browser: update 2012. Nucleic acids research, page gkr1012, 2011.

Pardis C Sabeti, David E Reich, John M Higgins, Haninah ZP Levine, Daniel J Richter, Stephen F Schaffner, Stacey B Gabriel, Jill V Platko, Nick J Patterson, Gavin J McDonald, et al. Detecting recent positive selection in the human genome from haplotype structure. Nature, 419(6909):832–837, 2002.

Pardis C Sabeti, Patrick Varilly, Ben Fry, Jason Lohmueller, Elizabeth Hostetter, Chris Cotsapas, Xiaohui Xie, Elizabeth H Byrne, Steven A McCarroll, Rachelle Gaudet, et al. Genome-wide detection and characterization of positive selection in human populations. Nature, 449(7164):913–918, 2007.

Tetsushi Sadakata and Teiichi Furuichi. Ca 2+-dependent activator protein for secretion 2 and autistic-like phenotypes. Neuroscience research, 67(3):197–202, 2010.

Rossana Sapiro, Igor Kostetskii, Patricia Olds-Clarke, George L Gerton, Glenn L Radice, and Jerome F Strauss III. Male infertility, impaired sperm motility, and hydrocephalus in mice deficient in sperm-associated antigen 6. Molecular and cellular biology, 22(17):6298–6305, 2002.

Carina M Schlebusch, Pontus Skoglund, Per Sjödin, Lucie M Gattepaille, Dena Hernandez, Flora Jay, Sen Li, Michael De Jongh, Andrew Singleton, Michael GB Blum, et al. Genomic variation in seven khoe-san groups reveals adaptation and complex african history. Science, 338(6105):374–379, 2012.

Andaine Seguin-Orlando, Thorfinn S Korneliussen, Martin Sikora, Anna-Sapfo Malaspinas, Andrea Manica, Ida Moltke, Anders Albrechtsen, Amy Ko, Ashot Margaryan, Vyacheslav Moiseyev, et al. Genomic structure in europeans dating back at least 36,200 years. Science, 346(6213):1113–1118, 2014.

Hifzur Rahman Siddique and Mohammad Saleem. Role of bmi1, a stem cell factor, in cancer recurrence and chemoresistance: preclinical and clinical evidences. Stem Cells, 30(3):372–378, 2012.

Adam Siepel, Gill Bejerano, Jakob S Pedersen, Angie S Hinrichs, Minmei Hou, Kate Rosenbloom, Hiram Clawson, John Spieth, LaDeana W Hillier, Stephen Richards, et al. Evolutionarily conserved elements in vertebrate, insect, worm, and yeast genomes. Genome research, 15(8):1034–1050, 2005.

John Maynard Smith and John Haigh. The hitch-hiking effect of a favourable gene. Genetical research, 23(01):23–35, 1974.

Karsten Suhre, Henri Wallaschofski, Johannes Raffler, Nele Friedrich, Robin Haring, Kathrin Michael, Christina Wasner, Alexander Krebs, Florian Kronenberg, David Chang, et al. A genome-wide association study of metabolic traits in human urine. Nature genetics, 43(6):565–569, 2011.

John A Todd, Neil M Walker, Jason D Cooper, Deborah J Smyth, Kate Downes, Vincent Plagnol, Rebecca Bailey, Sergey Nejentsev, Sarah F Field, Felicity Payne, et al. Robust associations of four new chromosome regions from genome-wide analyses of type 1 diabetes. Nature genetics, 39(7):857–864, 2007.

Ariel R Topletz, Jayne E Thatcher, Alex Zelter, Justin D Lutz, Suzanne Tay, Wendel L Nelson, and Nina Isoherranen. Comparison of the function and expression of cyp26a1 and cyp26b1, the two retinoic acid hydroxylases. Biochemical pharmacology, 83(1):149–163, 2012.

Meg Trahey, Gail Wong, Robert Halenbeck, Bonnee Rubinfeld, George A Martin, Martha Ladner, Christopher M Long, Walter J Crosier, Ken Watt, Kirston Koths, et al. Molecular cloning of two types of gap complementary dna from human placenta. Science, 242(4886):1697–1700, 1988.

ML Van Keuren, IM Hart, F-T Kao, RL Neve, GAP Bruns, DM Kurnit, and D Patterson. A somatic cell hybrid with a single human chromosome 22 corrects the defect in the cho mutant (ade-i) lacking adenylosuccinase activity. Cytogenetic and Genome Research, 44(2–3):142–147, 1987.

Cristiano Varin, Nancy Reid, and David Firth. An overview of composite likelihood methods. Statistica Sinica, 21(1):5–42, 2011.

Benjamin F Voight, Sridhar Kudaravalli, Xiaoquan Wen, and Jonathan K Pritchard. A map of recent positive selection in the human genome. PLoS biology, 4(3):e72, 2006.

Bruce S Weir, Lon R Cardon, Amy D Anderson, Dahlia M Nielsen, and William G Hill. Measures of human population structure show heterogeneity among genomic regions. Genome research, 15(11):1468–1476, 2005.

Danielle Welter, Jacqueline MacArthur, Joannella Morales, Tony Burdett, Peggy Hall, Heather Junkins, Alan Klemm, Paul Flicek, Teri Manolio, Lucia Hindorff, et al. The nhgri gwas catalog, a curated resource of snp-trait associations. Nucleic acids research, 42(D1):D1001–D1006, 2014.

Jay A White, Heather Ramshaw, Mohammed Taimi, Wayne Stangle, Anqi Zhang, Stephanie Everingham, Shelly Creighton, Shui-Pang Tam, Glenville Jones, and Martin Petkovich. Identification of the human cytochrome p450, p450rai-2, which is predominantly expressed in the adult cerebellum and is responsible for all-trans-retinoic acid metabolism. Proceedings of the National Academy of Sciences, 97(12):6403–6408, 2000.

Paul J Whiting, Timothy P Bonnert, Ruth M McKernan, Sophie Farrar, Beatrice Le Bourdelles, Robert P Heavens, David W Smith, Louise Hewson, Michael R Rigby, Dalip JS Sirinathsinghji, et al. Molecular and functional diversity of the expanding gaba-a receptor gene family. Annals of the New York Academy of Sciences, 868(1):645–653, 1999.

Yun-Yan Xiang, Shuhe Wang, Mingyao Liu, Jeremy A Hirota, Jingxin Li, William Ju, Yijun Fan, Margaret M Kelly, Bin Ye, Beverley Orser, et al. A gabaergic system in airway epithelium is essential for mucus overproduction in asthma. Nature medicine, 13(7):862–867, 2007.

Xin Yi, Yu Liang, Emilia Huerta-Sanchez, Xin Jin, Zha Xi Ping Cuo, John E Pool, Xun Xu, Hui Jiang, Nicolas Vinckenbosch, Thorfinnhorfinn Sand Korneliussen, et al. Sequencing of 50 human exomes reveals adaptation to high altitude. Science, 329(5987):75–78, 2010.

Alexandra Zhernakova, Clara C Elbers, Bart Ferwerda, Jihane Romanos, Gosia Trynka, Patrick C Dubois, Carolien GF De Kovel, Lude Franke, Marije Oosting, Donatella Barisani, et al. Evolutionary and functional analysis of celiac risk loci reveals sh2b3 as a protective factor against bacterial infection. The American Journal of Human Genetics, 86(6):970–977, 2010.

